# Random network structure stabilizes neural manifolds

**DOI:** 10.64898/2026.05.21.726949

**Authors:** Jens-Bastian Eppler, Santiago Galella, Gabriel C. Mel, Alex Roxin

**Affiliations:** Centre de Recerca Matemática, Campus de Bellaterra, Edifici C, 08193, Bellaterra, Barcelona, Spain; Frankfurt Institute for Advanced Studies, Ruth-Moufang-Str. 1, 60438, Frankfurt am Main, Germany; Barcelona Collaboratorium for Modelling and Predictive Biology, Dr. Aiguader 88, 08003, Barcelona, Spain

**Keywords:** Representational Drift, Neuronal Manifold Stability, Network Models, Artificial Neural Networks

## Abstract

Neuronal activity patterns change continuously over days and weeks, a phenomenon known as representational drift. Despite this, the geometric structure of population representations, namely the pairwise similarities between stimulus-evoked activity patterns, remains remarkably stable. How can ongoing changes in activity be consistent with stable representational similarity? We show that this is a generic consequence of random connectivity: in networks with random connectivity, output similarity is a monotonically increasing function of input similarity, independent of the specific connectivity pattern. Drift, whether driven by random synaptic turnover or Hebbian plasticity, merely transitions the network between random instantiations, leaving similarity intact. This extends to recurrent architectures and to deep neural networks, where continued training beyond performance saturation produces activity drift while preserving representational similarity. Although connectivity in the brain is not random, networks trained on high-dimensional inputs acquire connectivity that behaves statistically like a random projection, making these results broadly applicable to biological neural circuits.

## 1 Introduction

Representational drift is the phenomenon whereby neuronal activity patterns change over days and weeks, even in the absence of behavioral or environmental changes. It has been observed across a wide range of cortical and subcortical brain regions, including motor cortex (Rokni et al. 2007), barrel cortex (Margolis et al. 2012), parietal cortex (Driscoll et al. 2017), olfactory cortex (Schoonover et al. 2021), visual cortex (Deitch et al. 2021; Marks and Goard 2021; Bauer et al. 2024), auditory cortex (Aschauer et al. 2022), and hippocampus (Ziv et al. 2013; Geva et al. 2023), suggesting it is a general property of neuronal circuits (although notably reduced in some systems, such as the dentate gyrus (Hainmueller and Bartos 2018), bat hippocampus (Liberti et al. 2022), and the head-direction system of post-subiculum (Skromne Carrasco et al. 2026)). The most common theoretical frameworks cast representational drift as the natural consequence of ongoing learning. Even in the absence of explicit task demands Hebbian-like plasticity causes activity patterns to drift as each activity pattern will lead to changes in the network structure (Qin et al. 2023; Ratzon et al. 2024; Devalle et al. 2025; Eppler et al. 2026). Indeed, several reviews have argued that drift should come as no surprise: it is an inevitable byproduct of ongoing learning in any finite neural system, and in retrospect, existing theory could have predicted it all along (Duncker et al. 2020; Driscoll et al. 2022; O’Leary 2025). Recent work has extended this picture further, linking neuronal-level drift to systems consolidation via an entropic force (Kalle Kossio and Memmesheimer 2025), and to the ongoing extraction of environmental statistics through continuous statistical learning (Eppler et al. 2025). Complementary work has highlighted the role of stochastic synaptic changes - activity-independent fluctuations in synaptic strength arising from the inherent randomness of molecular processes at the synapse - as an additional driver of tuning instability (Yasumatsu et al. 2008; Loewenstein et al. 2011; Kasai et al. 2021). Together, these accounts provide a compelling picture of why drift occurs.

Despite the ongoing changes in neuronal activity patterns, behavior remains stable, day after day. How do unstable individual neurons give rise to stable behavior? One intriguing proposal is that neural circuits do not encode specific activity patterns, but rather the relationship between patterns: representation is, at its core, a representation of similarities (Shepard and Chipman 1970; Edelman 1998). Formalized as representational similarity analysis (Kriegeskorte et al. 2008; Kriegeskorte and Wei 2021), this perspective shifts attention from individual activity vectors to the geometry of population response vectors, often described as the low-dimensional manifold that stimulus-evoked activity traces out in neural state space (Gallego et al. 2017; Vyas et al. 2020; Perich et al. 2025). It is this geometry that turns out to be remarkably stable over time, even as the activity patterns that instantiate it drift. Across a growing number of brain regions and experimental paradigms, pairwise angles and correlations between population response vectors are preserved across sessions even as individual response vectors drift: in monkey motor cortex (Gallego et al. 2020), mouse hippocampus (Sheintuch et al. 2020; Levy et al. 2021; Keinath et al. 2022), and mouse auditory cortex (Noda et al. 2025). More recently, this has been characterized geometrically: drift in different brain regions can be described as a rigid transformation - a translation and rotation in population state space - that leaves the internal manifold geometry invariant (Aitken et al. 2022; Sylte et al. 2025; Eppler et al. 2025), a pattern arguably already visible in earlier observations (Ziv et al. 2013). What constrains drift to preserve neuronal manifold geometry remains an open question. Several mechanisms have been proposed: a stable anchor map or co-existing redundant maps of the same environment (Sheintuch et al. 2020), homeostatic plasticity (Noda et al. 2025), compensatory plasticity in downstream circuits (Kalle Kossio et al. 2021; Rule and O’Leary 2022), drift confined to a geometry-preserving subspace of equivalent solutions (Keinath et al. 2022; Sylte et al. 2025), and continuous statistical learning (Eppler et al. 2025). A similar dissociation between changing activity and preserved relational structure has also been observed in lateral entorhinal cortex (Kanter et al. 2025) and attributed there to an active circuit mechanism, as has been proposed in the hippocampal context as well (Keinath et al. 2022; Sylte et al. 2025). No consensus has emerged, and the central mechanistic question remains unresolved: What property of network connectivity and synaptic dynamics generically produces geometry-preserving drift?

Here we propose that a generic mathematical property of random connectivity, the preservation of pairwise similarities under random projections, provides a parsimonious account of manifold stability under drift. The core intuition is simple: when an output neuron receives input from many randomly connected upstream neurons, the similarity between its responses to two stimuli is determined by their overlap in input space, not by the specific pattern of connections. This is a statistical property of the ensemble, not of any particular instantiation, a consequence of the Johnson-Lindenstrauss lemma (Johnson and Lindenstrauss 1984) and related work on random projections in the context of compressed sensing and neural coding (Ganguli and Sompolinsky 2012). Because this holds for any sufficiently random connectivity with a large enough output population, it is robust to changes in specific wiring: synaptic turnover transitions the network from one random instantiation to another, leaving the pairwise similarity structure intact. Drift changes activity while largely preserving geometry.

The brain, however, is not random. We argue that this nevertheless applies, because any sufficiently large network trained on high-dimensional inputs will acquire connectivity that behaves statistically like a random projection. High-dimensional inputs impose strong constraints on network statistics that effectively prevent systematic exploitation of specific structure. Neural population activity in sensory cortex is indeed genuinely high-dimensional (Stringer et al. 2019), consistent with this regime. We show that the same similarity-preserving properties emerge in deep neural networks, where drift is modeled as continued training beyond performance saturation.

We demonstrate these similarity-preserving properties in a sequence of models of increasing biological realism. In binary feedforward networks with random connectivity, we show analytically and numerically that output similarity is a monotonically increasing function of input similarity, independent of the specific connectivity, and derive closed-form expressions for drift in activity and representational geometry under synaptic rewiring. This extends to Hebbian plasticity, recurrent architectures, and deep networks, where continued training beyond saturation produces activity drift while preserving representational similarity.

## 2 Results

### 2.1 Neuronal activity drifts while representational similarity is maintained

Representational drift, the ongoing change in neuronal activity patterns even under stable behavioral conditions, leaves the geometry of population responses, i.e. the pairwise similarities between stimulus-evoked activity vectors, largely intact (Ziv et al. 2013; Gallego et al. 2020; Noda et al. 2025). We illustrate this phenomenon in Fig. 1 with an example from mouse auditory cortex (raw data: Aschauer and Eppler (2022), processed data: Lai et al. (2025)). We pooled neuronal activity data across animals (18249 neurons total) and considered mean responses across trials to each of 19 pure tone stimuli ranging from 2.0 kHz to 45.0 kHz. Selecting the 500 most active cells and sorting them by preferred stimulus on imaging day 1 (see Methods for details), revealed the now typical signature of representational drift: the neuronal tuning curves span the entire range of stimuli, resulting in a clearly visible line along the ‘diagonal’ on day 1, that fades away with time (Fig. 1a). This effect is reversible, however: when selecting and sorting cells on the last imaging day, we see a time reversed pattern (Fig. 1b). These effects are not due to changes in the global population activity structure, since selecting and sorting neurons on each imaging day individually results in a stable pattern (Fig. 1c). The same effects are visible in the correlation of population response vectors (Fig. 1d, pooled across all neurons) and in the projection onto the first two dimensions using multi dimensional scaling (MDS, Fig. 1e, see Methods). While individual stimulus evoked activities are subject to change, the overall representational similarity structure is largely preserved.

**Fig. 1.**
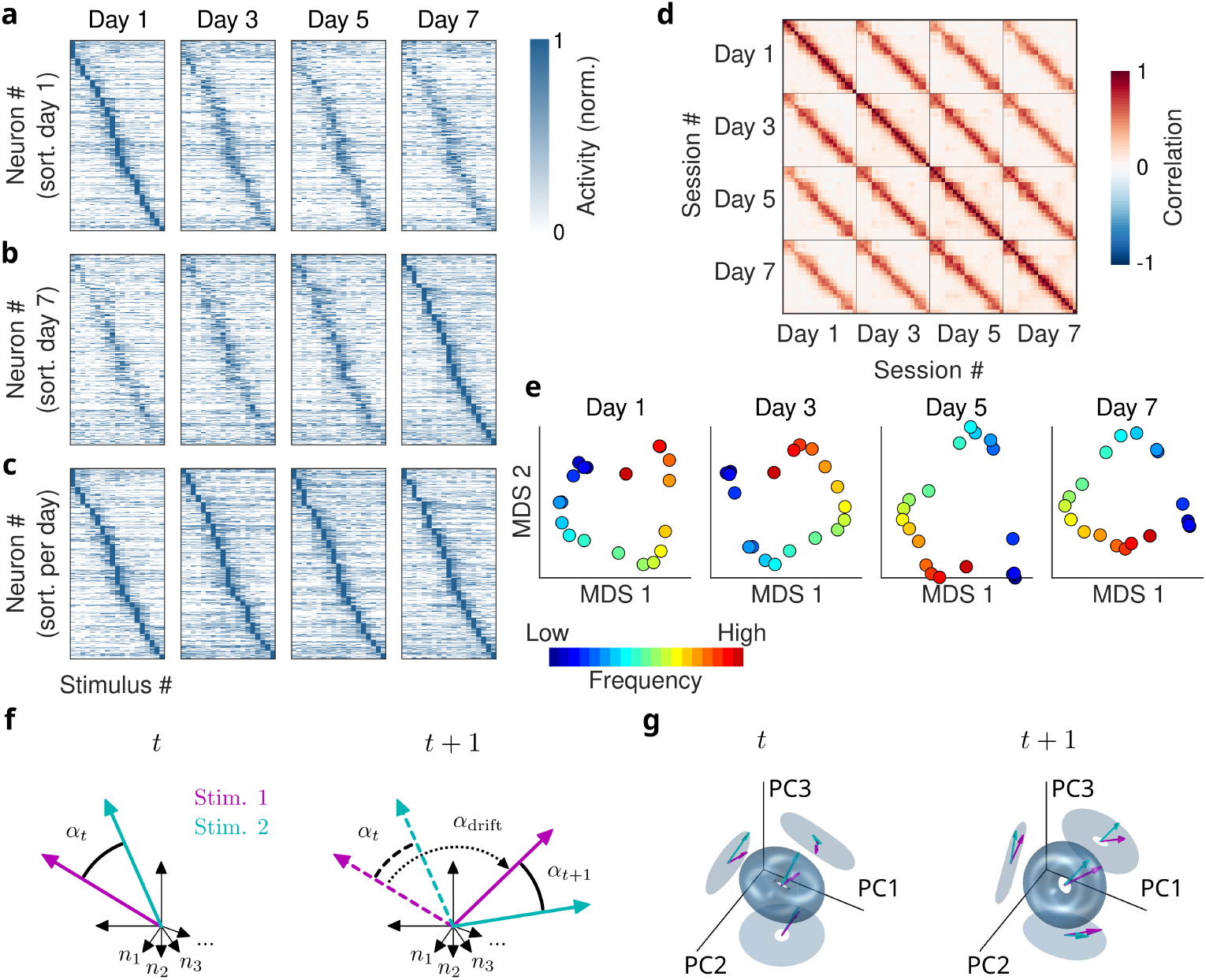
Representational drift changes activity patterns, but preserves representational similarities. **a)** Responses of auditory cortex neurons to pure tone stimuli, sorted by preferred stimulus on day 1. **b)** Same as (a), sorted on day 7. **c)** Same as (a, b), sorted on each day individually. **d)** Correlation matrix of responses in (a-c). **e)** Multi-Dimensional Scaling projection on the first two components per day. **f)** Schematic: Two stimuli evoke network responses with similarity *α*_*t*_ at time *t* (left). At *t* + 1 (right), responses drift by angle *α*_drift_ but maintain their similarity (*α*_*t*+1_ ≈ *α*_*t*_). **g)** Schematic: During representational drift, the similarity manifold (e.g., torus) rotates and may translate while preserving its structure. Magenta and cyan arrows show responses to two example stimuli.

On a given day, the representational similarity between two stimuli can be quantified by the angle *α*_*t*_ between their corresponding neural activity vectors (Fig. 1f, left). From one recording session to the next, individual response vectors drift by an angle *α*_drift_ (Fig. 1f, right). To distinguish changes in the neuronal activity from changes in representational similarity, we define two key metrics: *response vector drift*, which quantifies the change in neuronal response vectors to the same stimulus, and *response angle drift*, which measures the absolute change in the angle between response vectors and thus captures changes in representational similarity. Across recent findings (Sheintuch et al. 2020; Keinath et al. 2022; Noda et al. 2025), a consistent pattern has emerged: response vector drift substantially exceeds response angle drift, indicating that while individual neurons change their tuning properties, the neuronal activity manifold - the geometry of stimulus representations - remains largely intact. How can neuronal activity drift so dramatically while preserving the structure of the activity manifold? Fig. 1g illustrates a solution: if the representational manifold - here exemplified by a torus - undergoes rigid transformations such as rotations or translations, the relative positions of stimuli within the manifold remain fixed. Under such transformations, pairwise distances and angles are preserved, maintaining representational similarity although the neuronal activity patterns change. Crucially, any deformation beyond rigid transformations would distort these angles and thus alter the representational geometry. This raises the fundamental question: what synaptic changes in an underlying neural network can produce such drift?

### 2.2 In random networks, output similarity is a monotonic function of input similarity

To understand what network properties enable drift while preserving representational similarity, we first asked to what extent random feedforward networks can maintain the geometry of their input manifolds. We used a sparse binary network model

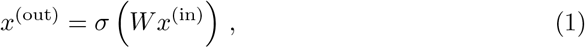

where *W* is a random binary connectivity matrix and *σ* is a threshold nonlinearity that maintains output sparsity (Fig. 2a, see Methods). We first tested the network with stimuli that have linearly dependent structure: neighboring stimuli are similar while distant stimuli are dissimilar, analogous to an animal navigating a linear track (Fig. 2b, see Methods). As expected, the network’s response vectors appear highly random and unstructured (Fig. 2c). However, when neurons are sorted by their preferred stimulus, a clear place field-like tuning emerges (Fig. 2d), revealing that the network preserves the linear structure of the inputs.

**Fig. 2.**
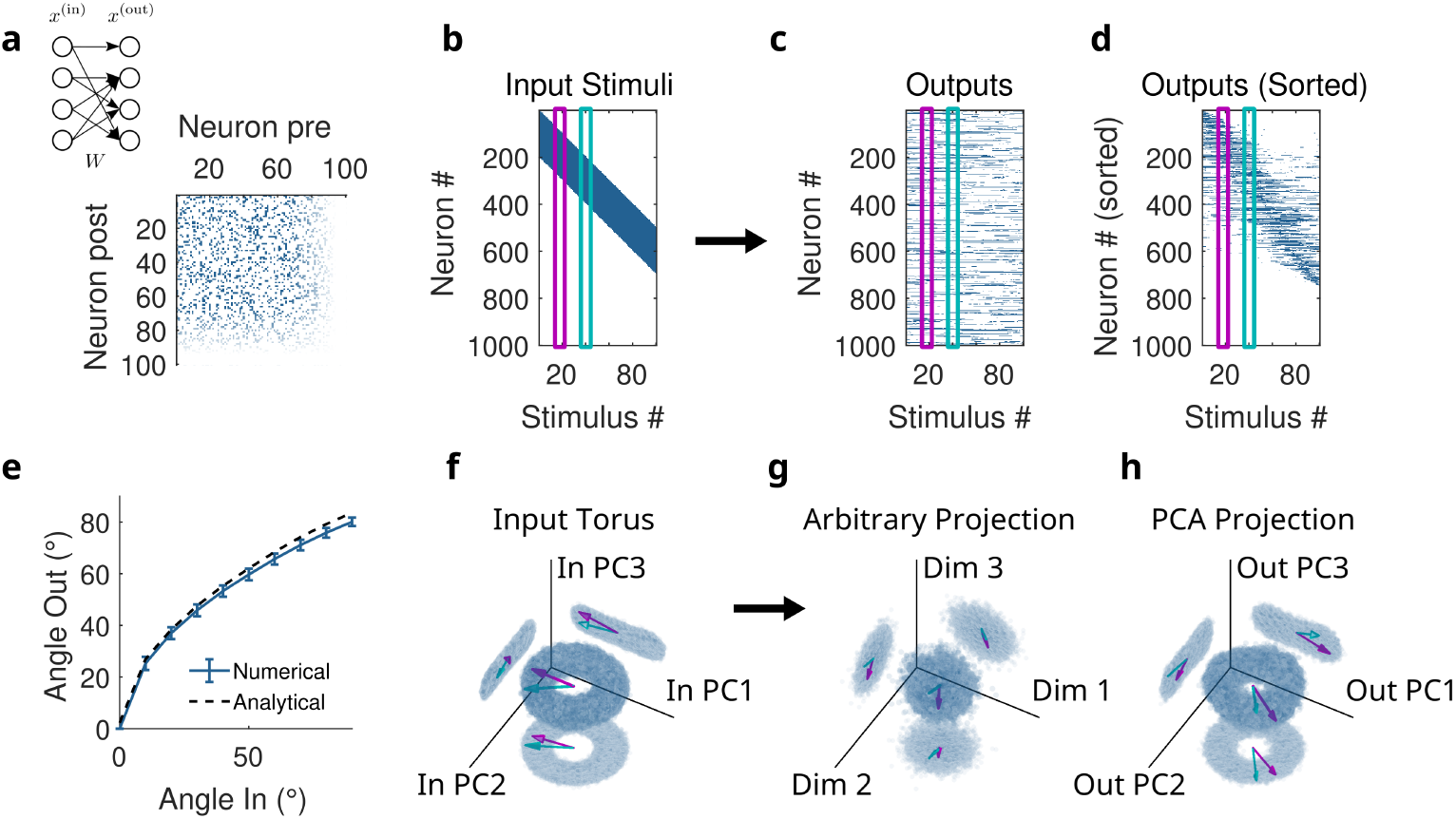
Random networks preserve input similarity relationships in their outputs. **a)** Feedforward network model (*x*^(out)^ = *σ*(*Wx*^(in)^)) with random binary connectivity matrix. *N* = 1000 neurons, sparsity 0.2. **b)** Input stimuli with linear structure, analogous to an animal on a linear track. Magenta and cyan highlight two example stimuli. **c)** Response vectors of the network in (a) to stimuli in (b). **d)** Same responses sorted by preferred stimulus. **e)** Output angle versus input angle. Solid line: simulations (mean ± SD, 1000 random networks, *N* = 1000, network sparsity 0.2, input sparsity 0.2). Dashed line: analytical solution (see Methods). **f)** First three principal components of toroidal inputs generated by random projection onto *N* = 1000 input neurons with binarization at sparsity 0.2 (see Methods). Cyan and magenta arrows show example stimuli. **g)** Projection of network responses (by network in (a) to stimuli in (f)) onto three random axes. Cyan and magenta indicate responses to example stimuli from (f). **h)** Projection of network responses (by network in (a) to stimuli in (f)) onto the first three principal components. Cyan and magenta indicate responses to example stimuli from (f).

To quantify this preservation, we systematically varied the input angle between stimulus pairs and plotted output angle as a function of input angle (Fig. 2e). The output angle is a monotonically increasing function of the input angle, indicating that the network faithfully preserves similarity relationships. We derived this input-output relationship analytically for this network by modeling pre-activations as bivariate Gaussian distributions and applying the threshold nonlinearity (see Methods). The analytical solution (dashed line) matches the numerical simulations (solid line).

This effect can also be seen with toroidal inputs. We generated toroidal stimuli by creating a torus in three dimensions, embedding it into high-dimensional space, and binarizing the result (Fig. 2f shows the first three principal components of these inputs, see Methods for details). Network responses to these inputs, when projected onto random axes (Fig. 2g) or onto the first three principal components of the outputs (Fig. 2h), reveal that the toroidal structure is maintained.

This preservation of input similarity is a generic consequence of random connectivity. When a neuron receives input from many randomly connected upstream neurons, the similarity between its responses to two stimuli is, on average, determined entirely by how similar those stimuli are, not by the particular pattern of connections. Because this holds for every output neuron independently of the specific weights, the population-level similarity structure of the inputs is reflected in the outputs. We derived this relationship analytically and confirmed that it matches simulations (see Methods for details).

Importantly, this preservation cannot be explained by randomly changing activity vectors alone, which would destroy the representational similarity structure for both linear and toroidal inputs (Supplementary Fig. 1).

### 2.3 Random synaptic changes cause activity drift while maintaining representational similarity

Since random networks generically preserve input similarity structure, we expected that random changes of the network connectivity would produce activity drift in network responses while largely leaving representational similarity intact. To test this, we simulated synaptic drift by randomly moving a fraction of synapses to new locations in the connectivity matrix (Fig. 3a). Using the same linear input stimuli as before (Fig. 3b), we tracked network responses across multiple sessions, moving 20% of synapses between each session.

**Fig. 3.**
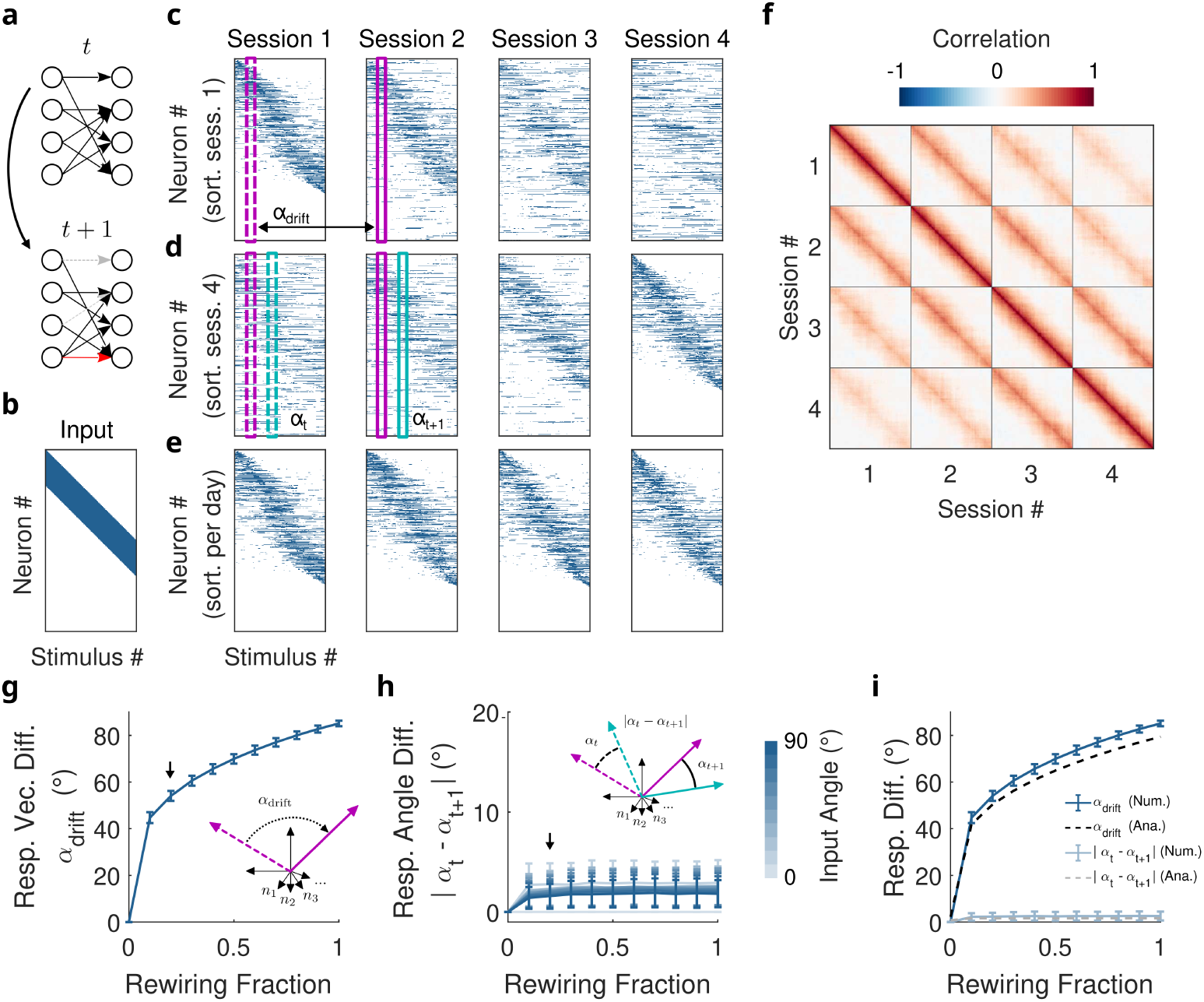
Random synaptic changes cause activity drift while preserving representational similarity. **a)** Schematic of random synaptic changes. **b)** Linear input stimuli (same as Fig. 2b). **c)** Network responses to stimuli in (b) across sessions. Between sessions, 20% of synapses are moved randomly. *N* = 1000, sparsity 0.2. Neurons sorted by preferred stimulus in session 1. **d)** Same as (c), neurons sorted by session 4 preference. **e)** Same as (c), neurons sorted by preference for each individual session. **f)** Correlation matrix of responses in (c)-(e). **g)** Response vector drift: change in network response to the same stimulus after randomly moving a fraction of synapses. Arrow indicates the session-to-session transition shown in (c). **h)** Response angle drift: absolute change in output angle between two responses (with given input angle) after randomly moving a fraction of synapses. Arrow indicates the session-to-session transition shown in (d). **i)** Response vector drift and response angle drift (at input angle 20°) versus fraction of randomly moved synapses. Solid line: simulations (mean ± SD, 1000 networks, network and input sparsity 0.2). Dashed line: analytical solution (see Methods).

When neurons are sorted by their preferred stimulus in session 1, we observe clear place field-like responses that progressively smear out over subsequent sessions (Fig. 3c), similar to experimental observations (Ziv et al. 2013; Aschauer et al. 2022). Critically, when we re-sort neurons by their preferred stimulus in the last session (Fig. 3d) or each session individually (Fig. 3e), the place field structure re-emerges, though with different neuronal identities. Neurons have changed their tuning, but overall the network maintains the representational similarity structure. This is also observable in the correlation matrix between network response vectors across stimuli and sessions (Fig. 3f). Within each session, we see the same linear pattern, fading away across sessions.

We quantified representational drift across sessions using the two metrics defined earlier: response vector drift, which measures changes in individual neuronal responses to the same stimulus (Fig. 3g), and response angle drift, which measures changes in representational similarity between stimulus pairs (Fig. 3h). The arrows mark the 20% synapse movement used in panels (c) to (e). Combining both metrics against the fraction of randomly moved synapses reveals that even small amounts of synaptic turnover produce substantial response vector drift, yet response angle drift remains low even when a large fraction of synapses have been moved (Fig. 3i). This demonstrates that random synaptic rewiring produces large activity changes while largely preserving representational similarity.

This preservation is again a consequence of random connectivity. When synapses are moved randomly, the network transitions from one random connectivity pattern to another. Because representational similarity is determined by the statistical properties of the connectivity rather than its specific realization, it remains stable across these transitions even as individual neural responses change substantially. We derived this relationship analytically, obtaining closed-form expressions for both response vector drift and response angle drift as a function of synaptic turnover (dashed lines Fig. 3i, see Methods). A key prediction of this analysis is that while response vector drift is independent of network size *N*, response angle drift decreases with *N* (Supplementary Fig. 2). This preservation follows from the Johnson-Lindenstrauss lemma (Johnson and Lindenstrauss 1984): random projections preserve pairwise distances with minimal distortion, even when projecting into lower dimensions. We observe, however, increasing variance in similarities with smaller output dimension (Supplementary Fig. 3). Notably, response angle drift, and thus the ratio of response angle drift to response vector drift, scales as 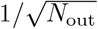 with output dimension *N*_out_, a relationship further illustrated in Supplementary Fig. 3i.

However, biological networks are not random. We next asked whether representational similarity is preserved when synaptic turnover is instead shaped by Hebbian plasticity.

### 2.4 Hebbian plasticity produces drift while maintaining representational similarity

We demonstrated that random synaptic changes largely preserve representational similarity during drift, but does this property extend to biologically plausible plasticity mechanisms? We examined Hebbian-like plasticity, a fundamental learning rule in which synaptic connections strengthen when pre- and post-synaptic neurons are co-active. We implemented a sparse binary Hebbian rule (Fig. 4a): when both pre- and post-synaptic neurons are active, a synapse forms with probability *p*_+_; when only one neuron is active, existing synapses are removed with probability *p*_−_; when both are inactive, the synapse remains unchanged. We varied *p*_+_ and *p*_−_ in parallel as a combined learning rate to explore the parameter space.

**Fig. 4.**
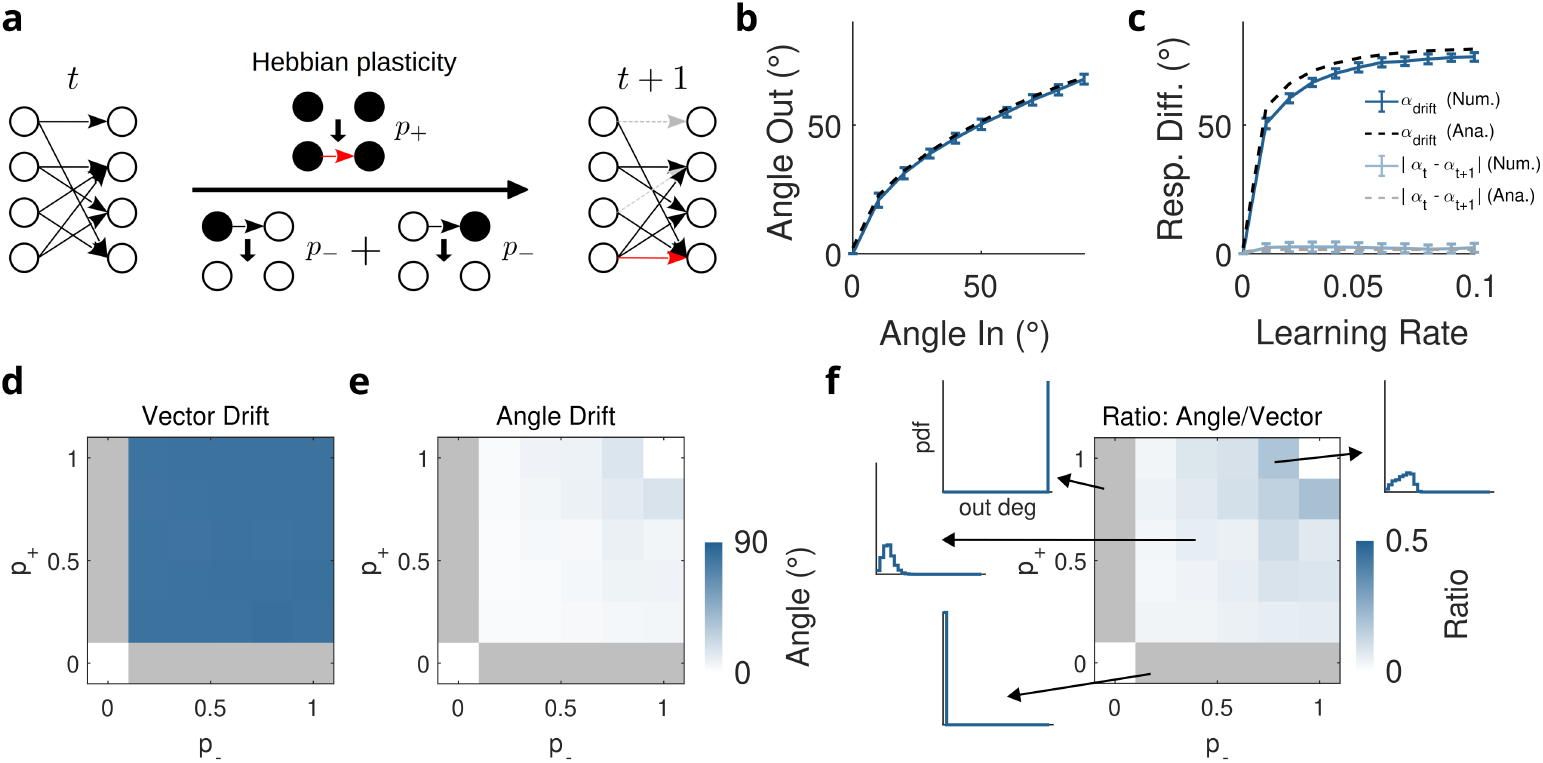
Hebbian plasticity causes activity drift while preserving representational similarity. **a)** Schematic of Hebbian plasticity. **b)** Output angle versus input angle. Solid line: simulations (mean ± SD, 100 networks, *N* = 1000, network sparsity 0.2, input sparsity 0.2, networks generated by 1000 Hebbian updates with random vectors and *p*_+_ = 0.1, *p*_−_ = 0.1). Dashed line: analytical solution (see Methods). **c)** Response vector drift and response angle drift (at input angle 20°) with Hebbian plasticity. Solid line: simulations (mean ± SD, 100 networks, sparsity 0.1, initialized as in (b), then 100 update steps with varying learning rates *p*_±_). Dashed line: analytical solution (see Methods). **d)** Response vector drift in Hebbian networks with varying *p*_+_ and *p*_−_. *N* = 1000, initialization as in (b), drift as in (c). Mean across 100 random networks. **e)** Response angle drift (input angle 20°) with varying *p*_+_ and *p*_−_. Same conditions as (d). **f)** Ratio of response angle drift (e) to response vector drift (d). Insets display the degree distribution of output connections.

To generate networks via Hebbian learning, we initialized random networks and applied 1000 Hebbian updates using sparse, random and binary pre- and post-synaptic activity patterns. We then tested whether these Hebbian-generated networks preserve input-output similarity. As with randomly generated networks (Fig. 2e), the output angle is a monotonically increasing function of input angle (Fig. 4b), demonstrating that Hebbian networks also maintain the geometry of their inputs. The analytical solution (dashed line, see Methods) matches the numerical simulations (solid line).

Starting from Hebbian-initialized networks, we applied 100 additional Hebbian update steps with varying learning strength *p*_±_ (Fig. 4c). As with random synaptic changes, response vector drift increases rapidly even with small amounts of plasticity, while response angle drift remains low even for strong plasticity. This demonstrates that Hebbian learning, like random synaptic turnover, produces substantial activity changes while maintaining representational similarity.

To explore the full parameter space, we varied the probabilities *p*_+_ and *p*_−_, separately, and measured both response vector drift (Fig. 4d) and response angle drift (Fig. 4e). Across all parameter combinations, response vector drift substantially exceeds response angle drift. This is quantified in the ratio plot (Fig. 4f): for most of the parameter space, the ratio remains well below 0.5, indicating strong preservation of representational similarity. Only when both *p*_+_ and *p*_−_ are very high does the ratio approach 0.5, indicating weaker preservation. The insets show the out-degree distribution for the different regimes. In this extreme regime, many inputs become mapped onto single outputs, decreasing the effective output dimension by introducing strong structure into the connectivity and disrupting the statistical properties that enable similarity preservation (see Supplementary Fig. 4 for complete in- and out-degree distributions across parameter space).

Importantly, similarity preservation also holds when Hebbian updates use correlated rather than random pre- and post-synaptic activity patterns. We tested four correlation structures: global correlations (each pattern correlated to one random “parent”), local correlations (each pattern correlated to the previous pattern), and combinations thereof (Supplementary Fig. 5). Across all correlation structures, response vector drift exceeds response angle drift, demonstrating that Hebbian plasticity preserves similarity even with structured, non-random activity patterns.

**Fig. 5.**
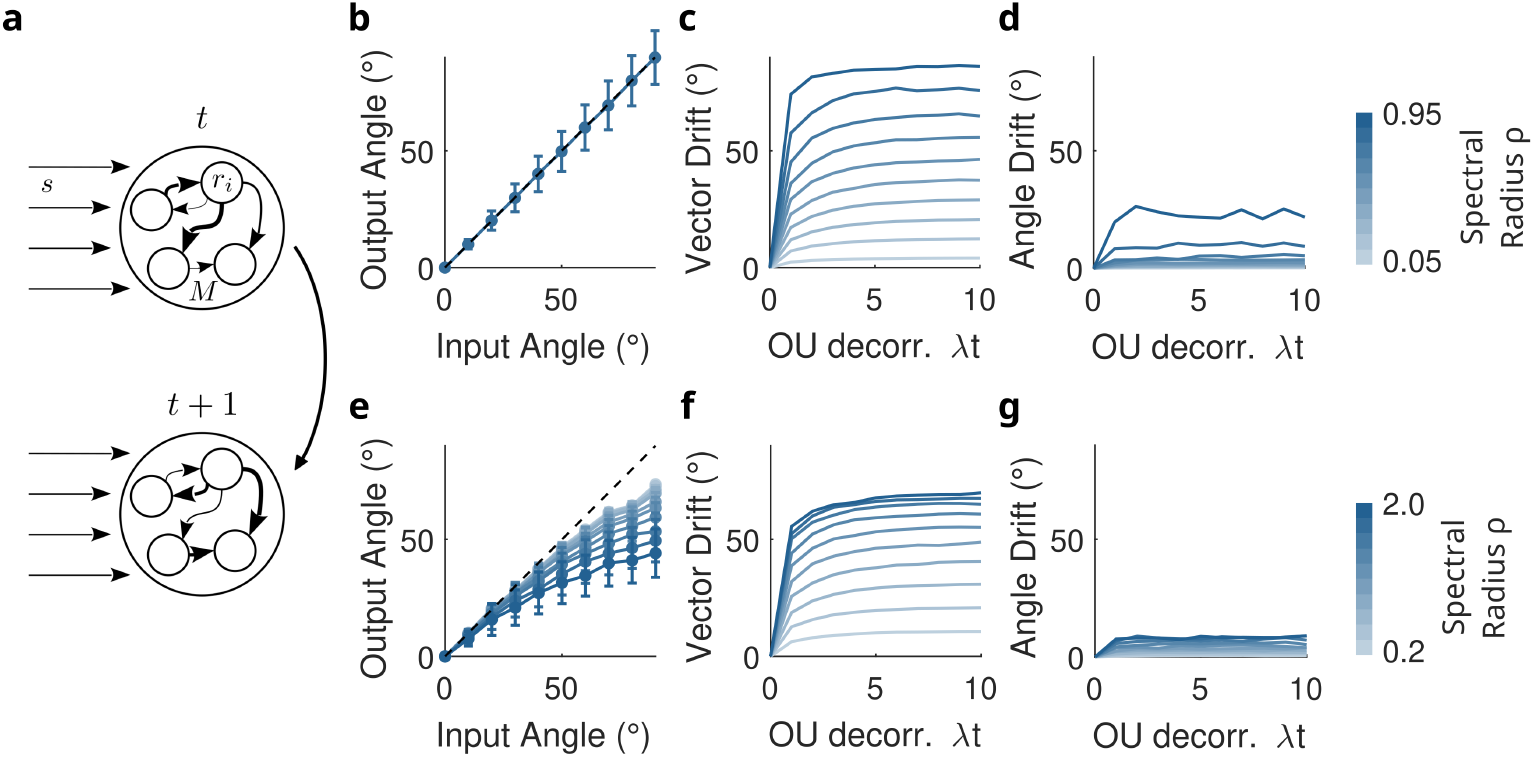
Random synaptic changes in recurrent networks cause activity drift while preserving representational similarity. **a)** Schematic of recurrent network with drift. **b)** Linear recurrent network preserves input angles: output angle = input angle. *N* = 200, Gaussian connectivity (*µ* = 0, 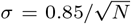), Gaussian inputs (*µ* = 0, *σ* = 1). Mean ± SD across 1000 networks. **c)** Response vector drift versus network drift, varying the spectral radius of the connectivity matrix. Network as in (b), drift via Ornstein-Uhlenbeck process. **d)** Response angle drift versus network drift, varying the spectral radius of the connectivity matrix. Network and drift as in (c). **e)** As (b) with half-wave rectification nonlinearity. **f)** As (c) with half-wave rectification nonlinearity. **g)** As (d) with half-wave rectification nonlinearity.

Finally, Hebbian plasticity can maintain biologically relevant features such as over-representation of particular stimuli. When certain stimuli are presented more frequently during Hebbian updates, using an asymmetric variant of the learning rule in which depression is gated by postsynaptic activity, they become selectively over-represented in the activity space (Supplementary Fig. 6, see Methods for details).

**Fig. 6.**
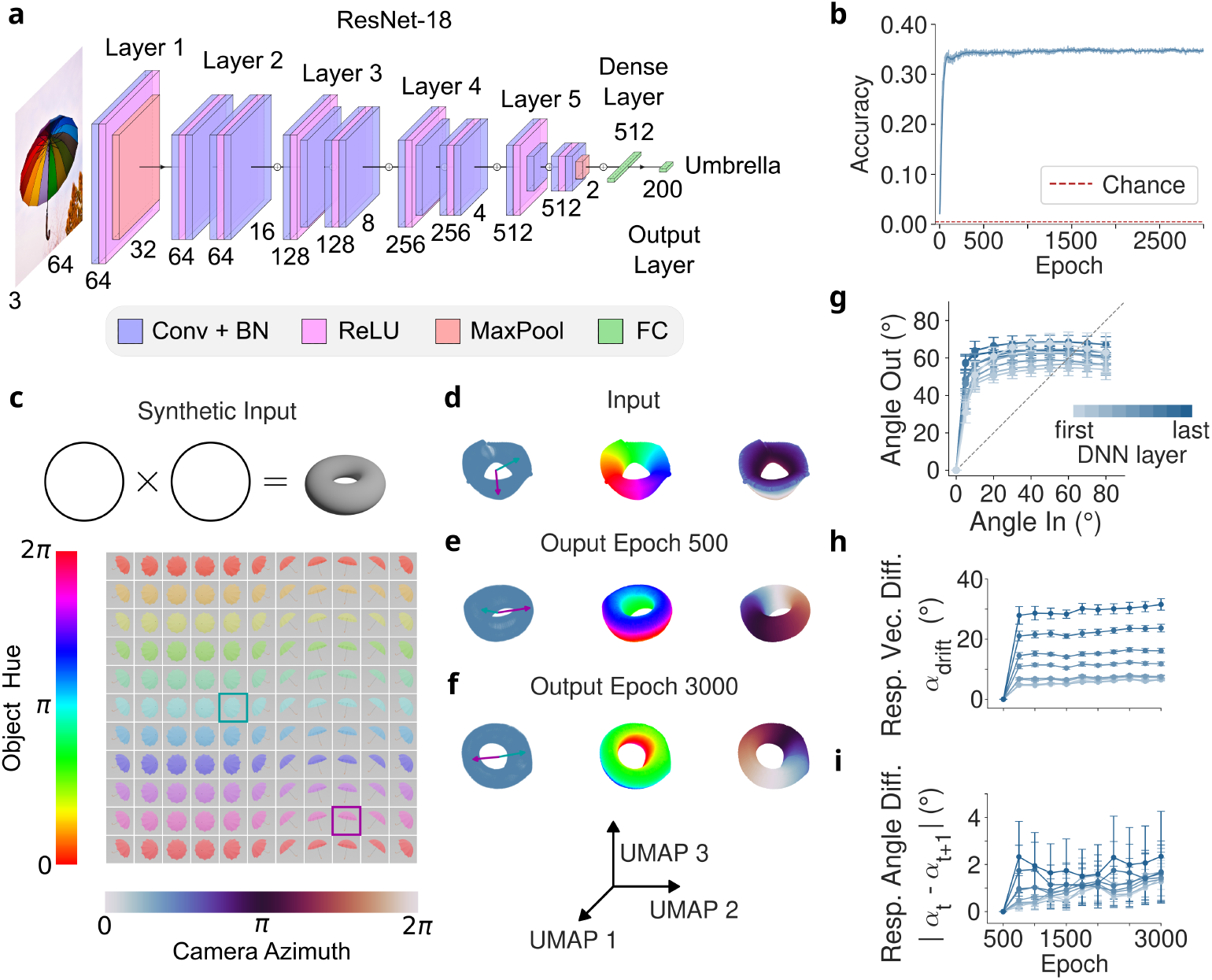
**a)** ResNet-18 model architecture. **b)** Training accuracy over epochs (validated on a held-out test set). **c)** Generation of a synthetic input with toroidal topology by sweeping object hue and camera azimuth for an umbrella object. **d)** UMAP projection of the input stimuli visualized in (c) using a grid of 100 rotation steps *×* 100 hue steps. **e)** UMAP projection of output of layer `layer2.relu1` at epoch 500 to the inputs in (d). **f)** UMAP projection of output of layer `layer2.relu1` at epoch 3000 to the inputs in (d). **g)** Input-output mapping across layers of the network. **h)** Response vector drift across layers of the network. **i)** Response angle drift across layers of the network.

This over-representation persists during continued drift, provided the presentation frequency is maintained. Thus, Hebbian plasticity can simultaneously produce drift, preserve representational similarity, and maintain task-relevant over-representation.

### 2.5 Random synaptic changes in recurrent networks cause activity drift while preserving representational similarity

We have demonstrated that both random synaptic changes and Hebbian plasticity largely preserve representational similarity in feedforward networks. However, many neural circuits are fundamentally recurrent. Do these preservation properties extend to recurrent architectures? To address this question, we switched from binary to Gaussian weights and inputs, as binary connectivity in rate-based recurrent networks is mathematically less tractable.

We began with linear recurrent networks, where the dynamics are

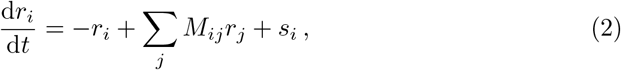

with Gaussian connectivity matrix *M* (Fig. 5a). In this linear case, mean output angles match input angles exactly (Fig. 5b), providing a trivial but important baseline in which geometry is preserved on average in the absence of nonlinearity.

We next examined how drift affects recurrent networks with different dynamical regimes by varying the spectral radius of the connectivity matrix. The spectral radius controls the strength of recurrent interactions: small values correspond to quasi-feedforward dynamics, while values close to 1 correspond to highly recurrent dynamics (for values > 1 network activity diverges). We implemented drift using an Ornstein-Uhlenbeck process (Uhlenbeck and Ornstein 1930), which maintains the Gaussian weight distribution in steady state. Across all spectral radii, response vector drift (Fig. 5c) substantially exceeds response angle drift (Fig. 5d), demonstrating that representational similarity is largely preserved across different recurrent regimes.

To make the model more biologically plausible, we introduced a half-wave rectification nonlinearity (ReLU), so the model reads

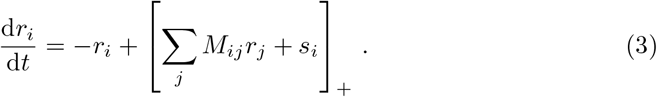

With this nonlinearity, output angles still closely track input angles (Fig. 5e), though the relationship deviates slightly from identity. Critically, the pattern of preservation remains: response vector drift (Fig. 5f) greatly exceeds response angle drift (Fig. 5g) across all spectral radii.

Thus, even in recurrent networks with biologically realistic nonlinearities, random synaptic changes produce substantial activity drift while maintaining representational similarity.

### 2.6 Input topology is reflected in deep neural network activities

Does the similarity-preserving property of random projections extend to trained deep neural networks, whose connectivity is far from random? To address this, we trained a ResNet-18 convolutional network (He et al. 2016) on Tiny ImageNet (Deng et al. 2009) and probed its internal representations using a structured set of stimuli with known geometry (Fig. 6a). The network was trained to classify natural images by mapping each input onto one of 200 discrete class labels (see Methods for details). ResNet-18 was chosen because the ResNet architecture is well-aligned with the primate ventral visual stream across datasets and tasks (Schrimpf et al. 2018; Zhuang et al. 2021; Xu and Vaziri-Pashkam 2021), making it a biologically relevant model system. Test set accuracy saturated by epoch 500 and remained stable throughout further training (Fig. 6b), which we used as our baseline.

To visualize the geometry of internal representations, we constructed a set of 10000 probe images from the MAPS dataset (Galella et al. 2026) by independently varying two circular stimulus dimensions, object hue and camera azimuth angle, on a regular grid, using a rendered umbrella object (Fig. 6c). Because both dimensions are circular, their combination defines a toroidal stimulus space, which is confirmed by a UMAP projection of the raw input images (Fig. 6d).

As shown by Galella et al. (2026) this toroidal topology is preserved in the network’s internal representations. At epoch 500, a UMAP projection of activations at an intermediate layer (`layer2.relu1`) reveals a clear toroidal structure (Fig. 6e): the network’s internal representation reflects the topology of the input space, despite being trained purely on object classification. This is further quantified by the input-output angle mapping across layers (Fig. 6g), which confirms that output similarity is a monotonically increasing function of input similarity. As argued above, this is expected: training on high-dimensional inputs drives the network’s connectivity into a regime that behaves statistically like a random projection, and similarity preservation follows to a large extent as a consequence.

### 2.7 Drift in deep neural networks maintains representational similarities

To assess the effects of representational drift in deep neural networks, we modelled drift as continued training beyond performance saturation (see Methods for details). This is in line with previous work showing that stochastic gradient descent noise alone drives ongoing changes in network representations after performance has stabilized (Pashakhanloo and Koulakov 2023; Ratzon et al. 2024). At epoch 3000, the toroidal structure remains clearly visible in the UMAP projection, but has undergone a rigid transformation - a rotation in the population activity space - relative to epoch 500 (Fig. 6f). This mirrors the geometry-preserving drift observed in biological circuits (Sylte et al. 2025; Eppler et al. 2025).

To quantify this, we computed response vector drift and response angle drift across layers and training epochs (Fig. 6h,i). All drift measurements represent changes relative to epoch 500, our reference point for comparison. As in the binary and recurrent network models, response vector drift substantially exceeds response angle drift throughout training, confirming that activity changes while representational similarity is preserved. Strikingly, both forms of drift increase with layer depth: later layers show greater activity changes and more distortion of the input-output angle mapping than earlier layers. This layer-dependent gradient mirrors observations in biology, where drift seems to be greater in higher-order regions (Aschauer et al. 2022; Ziv et al. 2013; Gallego et al. 2020). Together, these results demonstrate that similarity preservation under drift is not a special property of random networks, but extends naturally to trained deep neural networks, positioning them as a powerful model system for studying the mechanisms and consequences of representational drift.

## 3 Discussion

In this study, we showed that representational similarity is preserved as a generic mathematical consequence of random connectivity: in random networks, pairwise similarities between inputs are largely reflected in the outputs, independent of the specific connectivity pattern. Drift, whether random synaptic turnover or Hebbian plasticity, merely transitions the network between random instantiations, leaving this similarity intact. This property holds across architectures and extends to deep neural networks.

A direct consequence of this framework is that animals exposed to the same inputs will develop the same representational geometry - not because of shared developmental programs or co-ordinated plasticity, but simply as a mathematical consequence of the behavior of random projections. This parsimoniously explains the observed cross-animal consistency of neural representations within and across species (Rubin et al. 2019; Chen et al. 2021; Safaie et al. 2023; Codol et al. 2026), without invoking the additional mechanisms those studies required. A particularly vivid illustration comes from human color perception: we all perceive hue as lying on a circle, a geometry entirely determined by our three cone types serving as the input to downstream circuits (Shepard 1987, 1994; Brouwer and Heeger 2009; Rosenthal et al. 2021). You cannot know whether your blue is my blue, but we can agree on the distances between colors, and we do so not because our brains are wired alike, but because they receive the same input. Strikingly, this predicts that dichromats, whose inputs are two-dimensional rather than three, should perceive color not on a circle but along a line, a prediction consistent with what is known about dichromatic color space, and one that could be tested more rigorously with modern neural recording methods.

The brain, of course, is not random - and crucially, it can do more than passively reflect its inputs. Learning reshapes the representational manifold, selectively expanding or rotating distances between stimuli to support task-relevant distinctions (Sadtler et al. 2014; Aschauer et al. 2022; Flesch et al. 2022; Miguel-Lopez et al. 2025), a process also studied theoretically in recurrent networks (Driscoll et al. 2024). Within our framework, this raises a natural prediction: post-learning drift should gradually erode whatever structure learning imposed, pulling the manifold back toward the geometry dictated by the inputs alone. Yet behavior remains stable, so the learned representation must be anchored somewhere. Whether that anchor lies in primary sensory or motor cortex, in the hippocampus, or in higher cortical areas remains to be determined. At this point our parsimonious account reaches its limits, and active circuit mechanisms (Kanter et al. 2025; Eppler et al. 2025), as well as homeostatic plasticity (Noda et al. 2025) and compensatory changes in downstream circuits (Kalle Kossio et al. 2021; Rule and O’Leary 2022), become relevant candidates for how stability is ultimately enforced.

Our analytical results predict that geometry preservation improves with network size: response angle drift decreases with *N*, meaning that larger populations are better at preserving representational similarity under drift. This has a direct implication for experiments. When only a small fraction of neurons is recorded, the sampled population may give an impoverished picture of the true geometry: some stimuli may fail to evoke any response in the recorded neurons, and similar stimuli may evoke identical responses simply because the neuron that could distinguish them was not captured. This suggests that estimates of representational drift and geometry preservation from small recordings should be interpreted with caution, and that larger, denser recordings should reveal better-preserved geometry, a prediction that is now becoming testable with modern large-scale imaging methods.

Preserving the representational manifold does not solve the readout problem. A downstream neuron reading out via fixed synaptic weights would still be disrupted by drift, since the neurons carrying the representation change even as the geometry they instantiate does not. Some mechanisms have been proposed, compensatory plasticity in downstream circuits (Kalle Kossio et al. 2021; Rule and O’Leary 2022), but whether these generalize across systems and timescales remains unclear. This dissociation is sharpened by our findings in deep neural networks, where later layers show more drift than earlier ones, a pattern that mirrors the hierarchy observed in biology, from sensory cortex (Aschauer et al. 2022) through hippocampus (Ziv et al. 2013) to motor cortex (Gallego et al. 2020). This raises the question of how a stable behavioral output is read out from an increasingly drifting representation as one moves up the processing hierarchy, and investigating this with artificial neural networks, where the full network is accessible and drift can be precisely controlled, seems a particularly promising direction for future work.

Together, our results suggest a reframing: the stability of representational similarity in the presence of ongoing activity drift does not require special mechanisms; it is a mathematical consequence of the statistical structure of random projections, and thus a default property of large neural populations. What does require explanation is the departure from this default: how learning carves task-relevant structure into the manifold, how that structure is protected over time, and how behavior remains anchored to a drifting substrate. These are the questions our framework brings into sharper focus.

## 4 Methods

Here we provide an overview of the models and analyses used in this study. Detailed analytical derivations are provided in the Supplementary Information.

### 4.1 Experimental data and analyses (Fig. 1a-e)

We analyzed a publicly available chronic two-photon calcium imaging dataset from auditory cortex of awake, head-fixed mice, previously described in detail in Aschauer et al. (2022) and Eppler et al. (2026). Briefly, GCaMP6-expressing neurons in layers 2/3 were recorded across four imaging sessions separated by two days (80 fields of view, 10 mice, 18249 neurons total). During each session, 34 brief auditory stimuli (50-70 ms, 20 trials each) were presented in randomized order, comprising 19 pure tones (2-45 kHz) and 15 complex sounds. Raw and preprocessed data are publicly available at Aschauer and Eppler (2022) and Lai et al. (2025), respectively. For the analyses presented here, we restricted the dataset to the 19 pure tones. Single-neuron activity was defined as the mean Δ*F/F* in the 40 ms following stimulus onset, averaged across trials to yield one response vector per stimulus per session.

For visualisation, the 500 most active neurons (ranked by mean response across stimuli within a session) were selected and sorted by preferred stimulus. Activity maps were constructed using cells selected either from a fixed reference session (day 1 or day 7, held constant across panels) or independently for each session. Responses were normalised per neuron by dividing by that neuron’s maximum response in that session.

Representational similarity was assessed by computing Pearson correlation matrices across all neurons between every pair of stimuli, pooled across sessions.

To visualise the low-dimensional structure of stimulus representations, classical multidimensional scaling (MDS) was applied. A dissimilarity matrix was constructed as *D* = 1 − *ρ*, where *ρ* is the Pearson correlation between population response vectors. The matrix was double-centred and its two leading eigenvectors were retained, yielding a 2D embedding in which proximity reflects response similarity. Axis reflections were corrected across sessions by Procrustes alignment to the first session (reflections only).

### 4.2 Binary feedforward network model (Figs. 2, 3, Sup. Fig. 2,3)

We modelled a feedforward network of *N* = 1000 neurons with random binary connectivity. Each pair of neurons was connected with probability *g* = 0.2, yielding on average *K* = 200 presynaptic inputs per neuron. Network activity was governed by

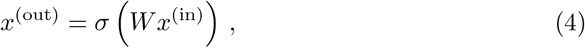

where *W* is the binary connectivity matrix and *σ* is a threshold nonlinearity: a neuron was active if its total input exceeded a fixed threshold *θ*. Input patterns were binary with sparsity *f* = 0.2. To preserve this sparsity at the output, *θ* was set analytically as the (1 − *f*)-th quantile of the Binomial(*K, f*) distribution, ensuring that the expected fraction of active output neurons matches that of the input.

To probe how output population size affects representational stability, we varied output dimension *N*_out_ ∈ {10, 100, 1000} while holding *N*_in_ = 1000 fixed.

### 4.3 Linear inputs (Figs. 2b, 3a, Sup. Fig. 1a)

To construct stimuli with a linear dependency structure, we generated 100 binary input patterns, each with sparsity *f* = 0.2 (200 active neurons out of *N* = 1000). Consecutive stimuli were shifted by a fixed step of *s* = (1 − *q*)*f N* neurons, where *q* = 0.975 is the pairwise overlap between successive stimuli, yielding a sliding window of active neurons that advances by 5 positions per step.

To visualise the output representations, neurons were sorted by their preferred stimulus, defined as the mean index of all stimuli eliciting a response, and ordered by this value in ascending order.

### 4.4 Toroidal inputs (Figs. 2f, Sup. Fig. 1c)

Toroidal stimuli were constructed by sampling a 100 × 100 parametric grid of points on a torus with major radius *R* = 3 and minor radius *r* = 1, yielding *N*_stim_ = 10000 points with coordinates (*x, y, z*) ∈ ℝ^3^. The resulting points were embedded in the full *N* -dimensional neural activity space by taking the (*x, y, z*) values to be the coordinates along 3 orthogonal axes in *N* -dimensional space. Axes were chosen by sampling 3 gaussian vectors and orthonormalising via QR factorisation. The continuous activation values were then passed through a sigmoid nonlinearity with a fixed temperature *T* = 0.7, and the offset was tuned by binary search to achieve a sparsity of *f* = 0.2. Input patterns were then obtained by probabilistic Bernoulli sampling of each neuron independently.

To visualise the geometry of the stimulus representations, principal component analysis (PCA) was applied to both the input and output populations. To assess how much of this structure is recoverable without knowledge of the principal axes, the output population activity was additionally projected onto three arbitrary orthonormal axes, again obtained by QR factorisation of a randomly drawn matrix.

### 4.5 Pairwise response similarity (Fig. 2e, Sup. Fig. 2a-c)

To characterise how the network transforms stimulus similarity, pairs of binary input patterns were constructed with a range of pairwise angles *θ*_in_ ∈ {0°, 10°, …, 90°*}*. Each stimulus had sparsity *f* = 0.2; the required overlap for a given angle was set as ⌊cos(*θ*_in_) · *fN*⌋, achieved by shifting a contiguous block of active neurons. The angle between the corresponding output patterns was computed as

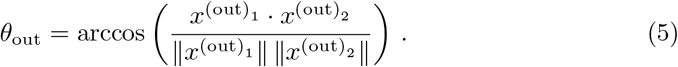

Numerical results were averaged over *n* = 1000 network realisations. Analytically, the expected output cosine similarity can be expressed as

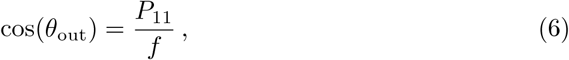

where *P*_11_ is the probability that both output neurons are simultaneously active, given by

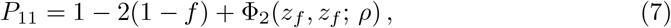

with *z*_*f*_ = Φ^−1^(1 − *f*), Φ_2_(·; *ρ*) the bivariate standard normal CDF with correlation *ρ*, and *ρ* the correlation of the pre-activation inputs (see Supplementary Methods for derivation).

### 4.6 Randomly changing response vectors (Sup. Fig. 1)

To verify that the sorting and dimensionality reduction pipelines do not artifactually impose structure on unstructured data, and to test that the observed representational structure is not robust to arbitrary random changes in the inputs, input stimuli were randomly perturbed prior to analysis. Each entry of the stimulus matrix was independently redrawn with probability *q* = 1 from a Bernoulli distribution with parameter *f*, such that all pairwise structure between stimuli is destroyed while the expected sparsity of each pattern is preserved. For the linear stimuli, the perturbed patterns were sorted by preferred stimulus using the same procedure as for the original inputs. For the toroidal stimuli, the perturbed patterns were projected onto three arbitrary orthonormal axes and submitted to PCA, again using the same pipeline as for the original inputs.

### 4.7 Binary network drift (Fig. 3)

To simulate gradual network change, synaptic drift was implemented by independently removing each existing connection with probability *d* and adding each absent connection with probability *dg/*(1 − *g*), where *g* = 0.2 is the connection probability and *d* is the rewiring fraction. This procedure preserves the expected sparsity of the network at all rewiring fractions *d* ∈ [0, 1], where *d* = 0 leaves the network unchanged and *d* = 1 redraws every connection independently. Numerical results were averaged over *n* = 1000 network realisations. Output activity for each stimulus and network realisation was computed as described above.

To visualise the output representations, neurons were sorted by preferred stimulus as described above, and pairwise Pearson correlation matrices were computed between all stimulus response vectors across the full population.

### 4.8 Response vector drift (Fig. 3g, Sup. Fig. 2d-f)

To quantify how much the response to a fixed stimulus changes as the network drifts, we computed the angle between the output vectors produced by the original and drifted network for the same input pattern,

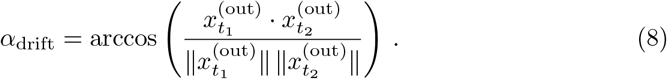

Numerical results were averaged over *n* = 1000 network realisations. Analytically, cos(*α*_drift_) is given by *P*_11_*/f* as above, where the pre-activation correlation is now determined by the overlap between the original and drifted connectivity matrix,

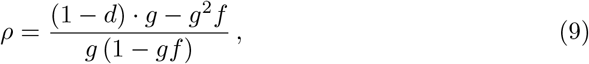

with *d* the rewiring fraction (see Supplemental Methods for derivation).

### 4.9 Response angle drift (Fig. 3h, Sup. Fig. 2g-i)

To quantify how much the angle between two stimulus responses changes as the network drifts, we computed

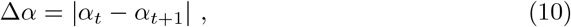

where *α*_*t*_ and *α*_*t*+1_ are the output angles between the same pair of stimuli measured on the original and drifted network, respectively. Numerical results were averaged over *n* = 1000 network realisations. Analytically, the expected signed difference ⟨*α*_*t*_ − *α*_*t*+1_ = 0⟩ by symmetry, so the expected absolute difference is determined entirely by the variance,

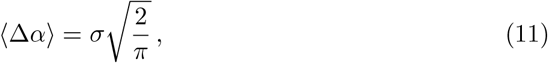

where *σ* is the standard deviation of the angle difference (see Supplemental Methods for derivation).

### 4.10 Hebbian network drift with uncorrelated inputs/outputs (Fig. 4, Sup. Figs. 4, 6)

Synaptic plasticity was implemented as a binary Hebbian learning rule (Devalle et al. 2025). At each update step, a pre-synaptic pattern *x*^(pre)^ and a post-synaptic pattern *x*^(post)^ were drawn independently from a Bernoulli distribution with sparsity *f*. Each synapse was then updated according to

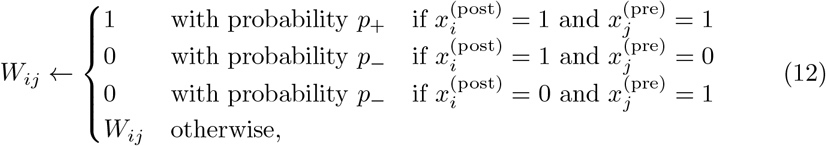

with *p*_+_ = *p*_−_ = 0.1. Networks were initialised with connection probability *g* = 0.2 and equilibrated over 1000 update steps, after which the expected connection probability converges to

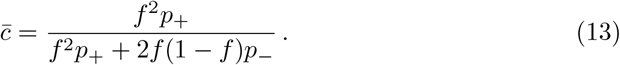

Synaptic drift was then simulated by continuing to apply the same learning rule for *T* = 100 additional steps, with scaled *p*_±_. The expected overlap between the original and drifted connectivity matrix decays as

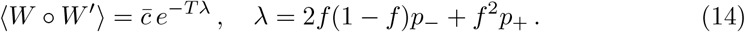

Numerical results were averaged over *n* = 1000 network realisations.

The analytical expressions for the input-output mapping, response vector drift, and response angle drift were derived within the bivariate normal framework described above, with *ρ* now corrected for the correlation structure of the Hebbian-equilibrated network. Specifically, while for a random binary network connections are independent, the Hebbian rule introduces correlations between synapses onto the same postsynaptic neuron, characterised by the joint potentiation probability 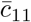 (Devalle et al. 2025). The pre-activation correlation *ρ* was derived accordingly, replacing the independent-synapse assumption with expressions involving 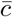 and 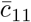 (see Supplementary Methods for derivation). The dependence of in- and out-degree distributions on *p*_+_ and *p*_−_ was characterised by sweeping both parameters independently over [0, 1] in steps of 0.2.

For the over-representation analyses, we used an asymmetric variant of the learning rule in which depression occurs only when the postsynaptic neuron is active and the presynaptic neuron is inactive, i.e. the third case in Eq. (12) is replaced by *W*_*ij*_ ← *W*_*ij*_. This asymmetry is necessary for over-representation to emerge: with the symmetric rule, a frequently-presented stimulus actively depresses connections onto neurons that do not respond to it, preventing the selective strengthening of a specific subpopulation. With postsynaptic gating of depression, by contrast, only neurons that already respond to the frequent stimulus undergo plasticity, allowing a positive feedback that builds up over-representation selectively. All other parameters and procedures were identical to those described above.

### 4.11 Hebbian network drift with correlated inputs/outputs

To investigate how correlations in the learning patterns affect synaptic drift, we repeated the analysis with structured pre- and post-synaptic patterns. Patterns were generated in one of two modes. In the *global* mode, each pattern was drawn by sampling *f N* active neurons such that a fraction *ρ* of them were shared with a fixed parent pattern, while the remaining active neurons were drawn from those inactive in the parent. In the *local* mode, each new pattern was drawn in the same way, but using the *previous* pattern as the parent, so that correlations were between successive patterns rather than to a common reference. For *ρ >* 0 the patterns are positively correlated, for *ρ* = 0 they are independent, and for *ρ <* 0 active neurons are preferentially drawn from those inactive in the parent.

Pre- and post-synaptic patterns were generated independently, each in either global or local mode, yielding four conditions: global pre/global post, global pre/local post, local pre/global post, and local pre/local post. For each condition, the pre-synaptic correlation *ρ*_pre_ and post-synaptic correlation *ρ*_post_ were swept independently over {−1, −0.5, 0, 0.5, 1}, giving a 5 × 5 grid of parameter combinations. Networks were initialised with connection probability *g* = 0.2, equilibrated over 1000 update steps using the symmetric Hebbian rule, and then drifted for 100 additional steps. Response vector drift and response angle drift were computed as described above. Results were averaged over *n* = 10 network realisations per condition.

### 4.12 Recurrent network drift (Fig. 5)

We compared the input-output mapping and drift properties of linear and nonlinear recurrent networks. In both cases, the connectivity matrix *M* was initialised as a random Gaussian matrix with zero diagonal, scaled to a target spectral radius λ_max_. Input stimuli were unit-norm Gaussian vectors with pairwise angles set exactly via Gram-Schmidt orthogonalisation.

For the linear network, the steady-state response to an input *x* was given by

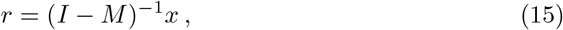

which is defined for λ_max_ *<* 1. For the nonlinear network, the response was obtained by integrating the dynamics

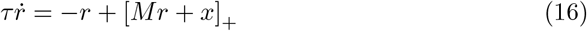

to steady state using forward Euler with time step Δ*t* = 0.01 and settling time 5*τ*, where *τ* = 1 and [·]_+_ denotes rectification.

Synaptic drift was implemented as an Ornstein-Uhlenbeck (OU) process on the connectivity matrix (Uhlenbeck and Ornstein 1930). At each time step *t*, the network evolved as

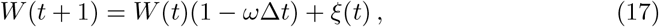

where *ω* = 0.3 is the OU decay rate and *ξ*(*t*) is a matrix of independent Gaussian noise terms with variance 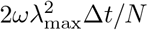, scaled to maintain the spectral radius in expectation. The diagonal of *M* was set to zero after each update. Responses and drift were evaluated at *n*_*t*_ = 11 time points separated by Δ*t* = 1, and averaged over *n* = 100 network realisations. The spectral radius was varied over λ_max_ ∈ [0.05, 0.95] for the linear network and λ_max_ ∈ [0.2, 2.0] for the nonlinear network.

### 4.13 ResNet-18 training and drift protocol (Fig. 6)

We trained a ResNet-18 (He et al. 2016), comprising 17 convolutional layers with batch normalization and ReLU activations followed by a single fully connected layer, on the Tiny ImageNet dataset (Deng et al. 2009), which consists of 100,000 images of resolution 64 × 64 pixels spanning 200 object classes. During training, we employed a standard suite of augmentations including random resized crops (64×64 resolution, scale [0.8,1.0]), random horizontal flips (p=0.5), and color jittering (brightness/contrast/saturation factor of 0.2; hue factor of 0.02). We also applied a 10% probability of grayscale conversion. All inputs were normalized using training-set mean and standard deviation values prior to being fed into the network. Optimization was performed using stochastic gradient descent with a fixed learning rate of 0.01, a batch size of 256, with weight decay of 1e-4 and no momentum. The network was trained for 3000 epochs using cross-entropy loss with model weights checkpointed every 10 epochs. Performance on the test set stabilized by epoch 500 (test accuracy 34. ± 1.33%, peak-to-peak variation 2.33% from epoch 500 onward). We defined epoch 500 as the baseline and treated continued training beyond this point as our model of representational drift, in line with previous work showing that stochastic gradient descent noise alone is sufficient to drive drift in trained networks (Pashakhanloo and Koulakov 2023; Ratzon et al. 2024).

### 4.14 Structured probe stimuli (Fig. 6)

To probe the geometry of internal representations, we generated a structured set of 10,000 images using the MAPS dataset (Galella et al. 2026). An umbrella object (a class present in Tiny ImageNet) was rendered using Blender with the Cycles renderer (Blender Online Community 2025), sweeping 100 values of camera azimuth angle and 100 values of object hue, sampled on a regular grid. Both dimensions are circular, so their combination defines a toroidal stimulus space. The object’s asymmetry ensured that distinct azimuth angles produced distinct activation patterns.

### 4.15 Layer-wise activation analysis (Fig. 6)

We extracted activations from nine layers of the trained ResNet-18, corresponding to the output to the ReLU activation function: `layer1.relu1`, `layer2.relu1`, `layer2.relu2`, `layer3.relu1`, `layer3.relu2`, `layer4.relu1`, `layer4.relu2`, `layer5.relu1`, `layer5.relu2`. Activations were spatially flattened to vectors prior to analysis. Representational geometry was characterized using the same response vector drift and response angle drift metrics defined above. To visualize the low-dimensional structure of representations, we first applied PCA retaining 20 components, followed by UMAP (McInnes et al. 2018) projected onto 3 dimensions.

## Supplementary information

This article is accompanied by supplementary material, including Supplementary Figures 1–5 and detailed analytical derivations in the Supplementary Methods.

## Acknowledgements

We thank members of the Computational Neuroscience group at the Centre de Recerca Matemàtica and the BARCCSYN community for great feedback throughout this project. We are grateful to Simon Rumpel and Johannes Seiler for helpful discussions. The auditory cortex data used in Fig. 1 were collected by Dominik Aschauer.

## Data availability

The raw and processed imaging data used in this study are publicly available at https://doi.org/10.12751/g-node.trwj8c and https://doi.org/10.12751/g-node.vnj8ip, respectively.

## Code availability

Code to reproduce all simulations and analyses will be made available upon publication.

## Author contribution

JBE and AR conceived the study. JBE, SG and GM developed the methodology and performed formal analysis. JBE and SG developed the software, carried out the investigation, and validated the results. GM contributed to the investigation. JBE and SG created the visualizations. JBE wrote the original draft. JBE, SG, GM and AR reviewed and edited the manuscript. AR supervised the project and acquired funding.

