## Supplementary Information for "Random network structure stabilizes neural manifolds"

for

**Supplementary Methods** **9**

**Supplementary References** **14**

### Supplementary Figures

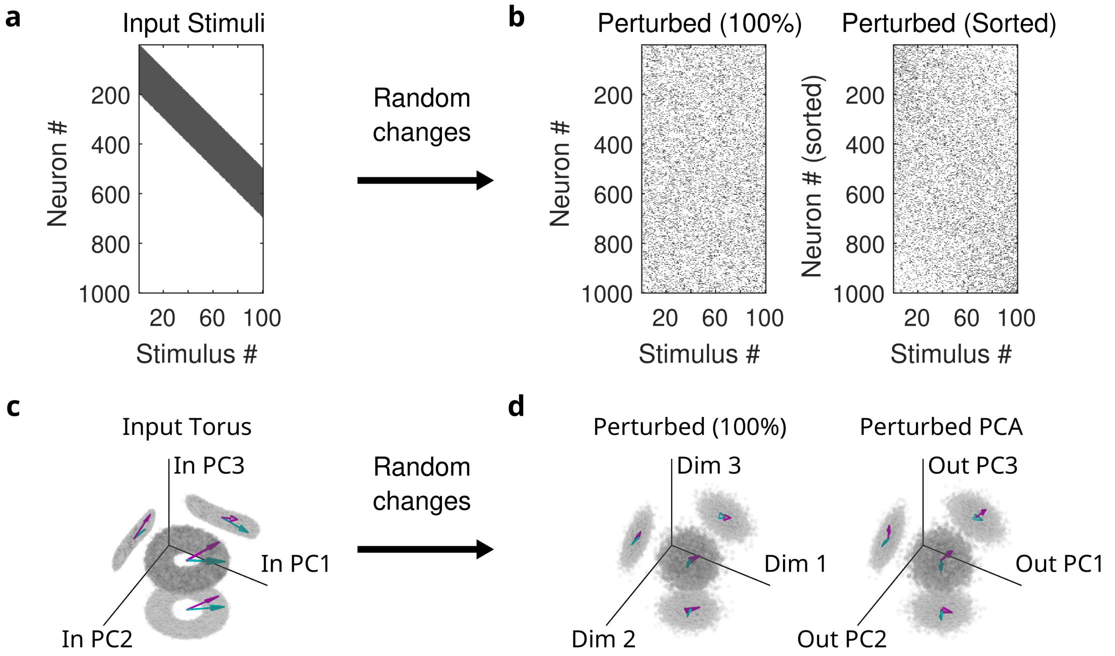

**Supplementary Fig. 1: Randomly changing vectors does not preserve representational similarity. a)** Input stimuli with linear structure, analogous to an animal on a linear track. **b)** Randomly changing activity vectors does not preserve input structure. Left: same sorting as in (a). Right: Sorting by preferred stimulus. **c)** First three principal components of toroidal inputs generated by random projection onto $N = 1000$ input neurons with binarization at sparsity 0.2 (see Methods). Cyan and magenta arrows show example stimuli. **d)** Randomly changing activity vectors does not preserve input structure. Left: projection onto three random axes. Right: Projection onto the first three principal components. Cyan and magenta indicate responses to example stimuli from (c).

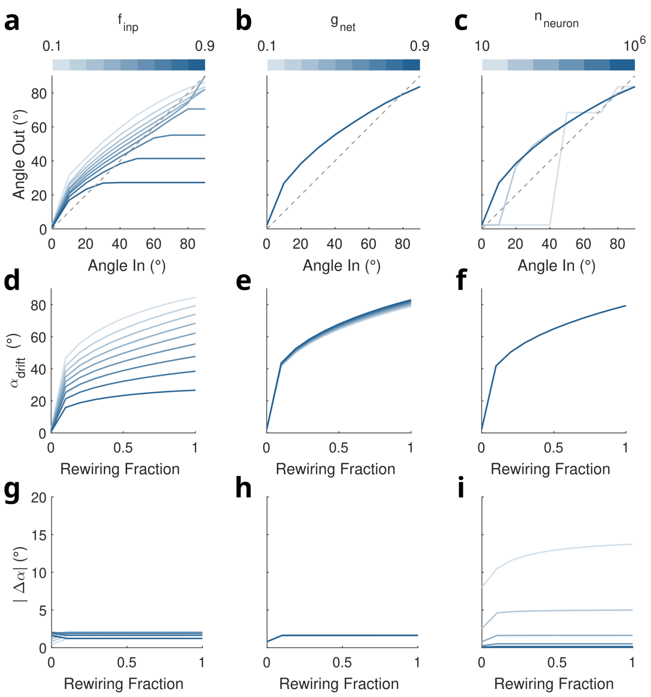

**Supplementary Fig. 2: Effects of input sparsity, network sparsity and network size on input-output mapping, response vector drift and response angle drift.** Varying input sparsity $f_{\mathrm{inp}}$, network sparsity $g_{\mathrm{net}}$ and network size $n_{\mathrm{neuron}}$ reveals stability of the observed effects. While systematically varying one of the three the others were kept at $f_{\mathrm{inp}} = 0.2$, $g_{\mathrm{net}} = 0.2$ and $n_{\mathrm{neuron}} = 1000$. **a)** Input-output mapping varying input sparsity $f_{\mathrm{inp}}$. **b)** Input-output mapping varying network sparsity $g_{\mathrm{net}}$. **c)** Input-output mapping varying network size $n_{\mathrm{neuron}}$. **d)** Response vector drift varying input sparsity $f_{\mathrm{inp}}$. **e)** Input-output mapping varying network sparsity $g_{\mathrm{net}}$. **f)** Input-output mapping varying network size $n_{\mathrm{neuron}}$. **g)** Response angle drft varying input sparsity $f_{\mathrm{inp}}$. **h)** Input-output mapping varying network sparsity $g_{\mathrm{net}}$. **i)** Input-output mapping varying network size $n_{\mathrm{neuron}}$. Note that while response vector drift is high and independent of network size (f), response angle drift decreases with growing network size (i).

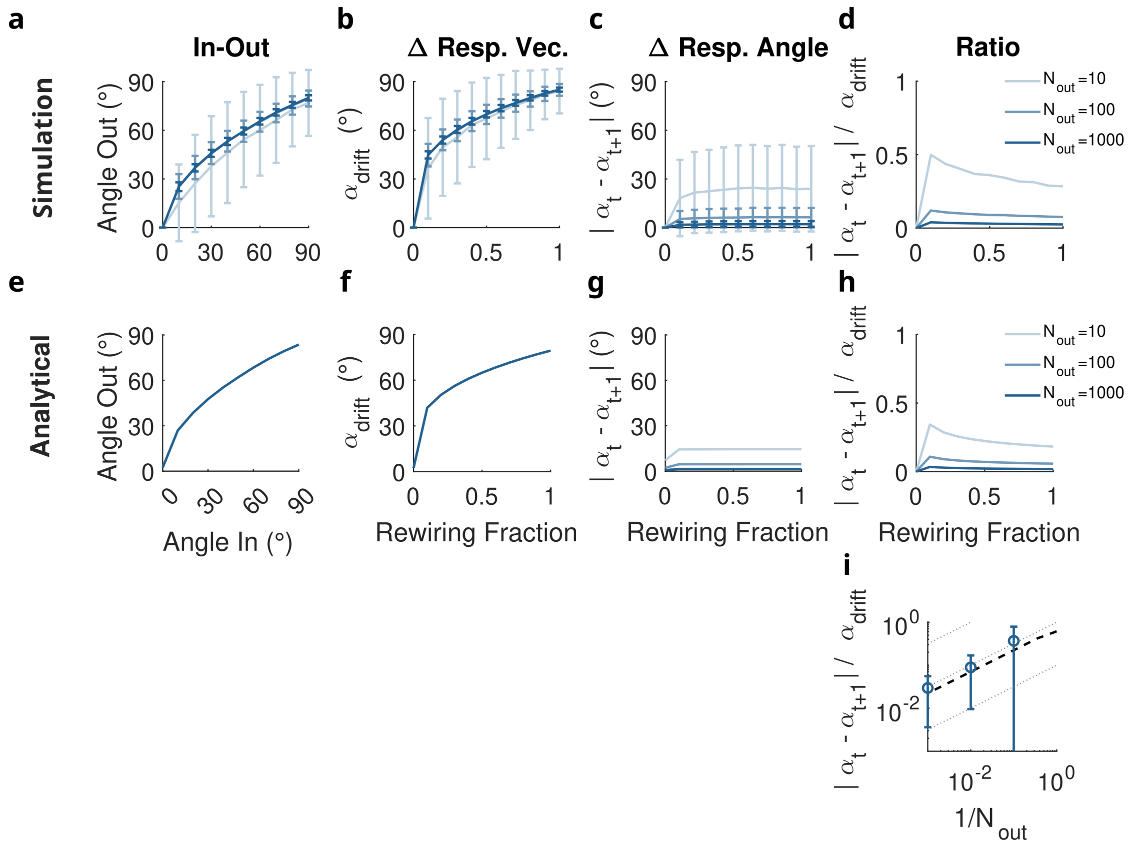

**Supplementary Fig. 3: Output dimensionality modulates the fidelity of representational similarity preservation under drift.** While input dimensionality is held constant at $N_{\text{in}} = 1000$, output dimensionality $N_{\text{out}}$ is varied across $\{10, 100, 1000\}$. Network sparsity $g = 0.2$ and input sparsity $f = 0.2$ are fixed throughout. **a)** Input-output mapping (numerical). **b)** Response vector drift (numerical). **c)** Response angle drift (numerical). **d)** Ratio of response angle drift to response vector drift (numerical). **e)** Input-output mapping (analytical). **f)** Response vector drift (analytical). **g)** Response angle drift (analytical). **h)** Ratio of response angle drift to response vector drift (analytical). **i)** Ratio of response angle drift to response vector drift vs. the output dimensionality for a fixed rewiring fraction of 0.5 shows scaling with $1/\sqrt{N_{\text{out}}}$. Blue: numerical mean ± SD across 1000 networks. Dashed: analytical. Grey lines: $f(x) = a\sqrt{x}$ with $a = \{.1, 1, 10\}$.

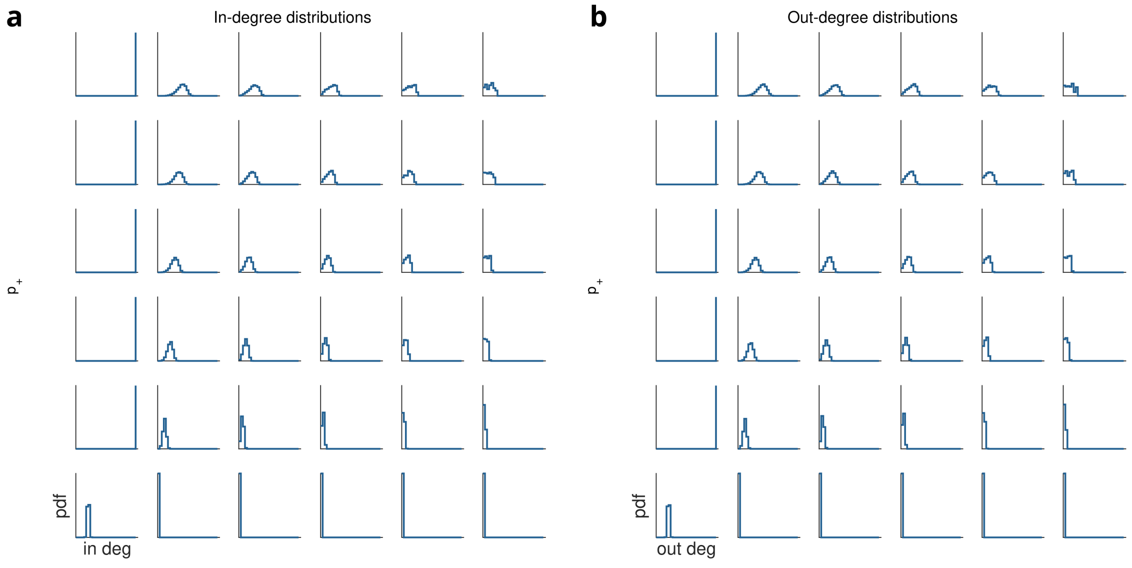

**Supplementary Fig. 4: In- and out-degree distributions for varying Hebbian parameters $p_+$ and $p_-$.** **a)** In-degree distribution with varying $p_+$ and $p_-$ (both from 0 to 1). When $p_+ = 0$, the distribution is a delta peak at 0; when $p_- = 0$, it is a delta peak at 1. When both $p_+ = p_- = 0$, no drift occurs. For other parameter combinations, the distribution broadens substantially, becoming increasingly wide for large $p_+/p_-$. **b)** Out-degree distribution with varying $p_+$ and $p_-$ (both from 0 to 1). When $p_+ = 0$, the distribution is a delta peak at 0; when $p_- = 0$, it is a delta peak at 1. When both $p_+ = p_- = 0$, no drift occurs. For other parameter combinations, the distribution broadens substantially, becoming increasingly wide for large $p_+/p_-$.

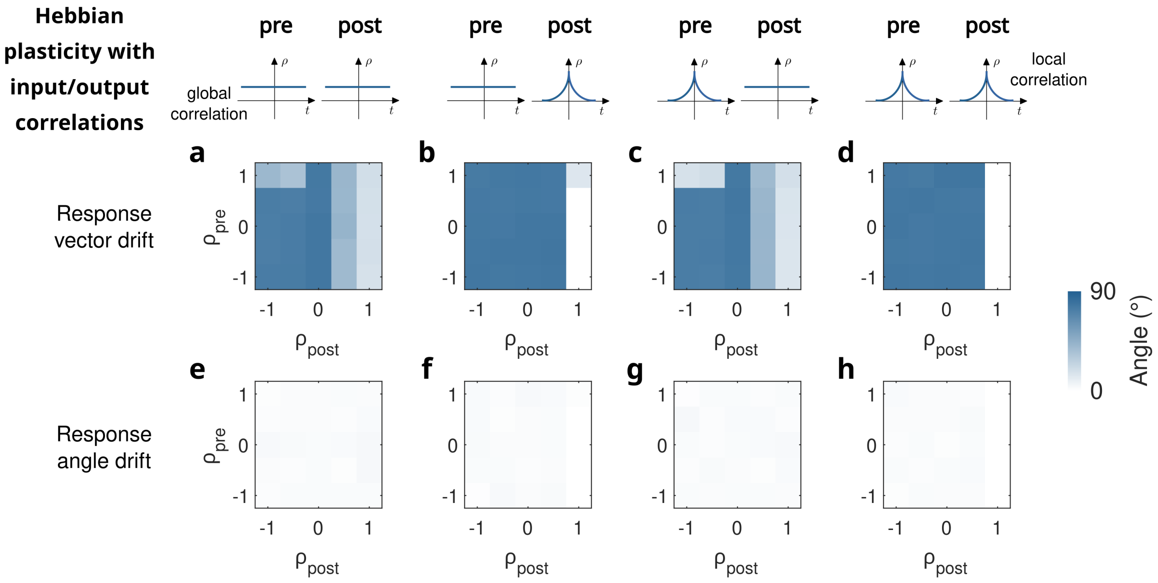

**Supplementary Fig. 5: Correlated Hebbian updates preserve representational similarity despite response drift. a)-d)** Response vector drift with correlated pre- and post-synaptic activity in Hebbian updates. $N = 1000$, network sparsity 0.2, stimulus sparsity 0.2, input/output sparsity 0.2, $p_+ = p_- = 0.1$, $\eta = 1$, initialization and drift as in Fig. 4. Mean over 10 networks. Correlations were either global (each pattern correlated to one random "parent") or local (each pattern correlated to previous pattern, see Methods). **a)** Global pre and post. **b)** Global pre, local post. **c)** Local pre, global post. **d)** Local pre and post. **e)-h)** Same as (a)-(d) but showing response angle drift.

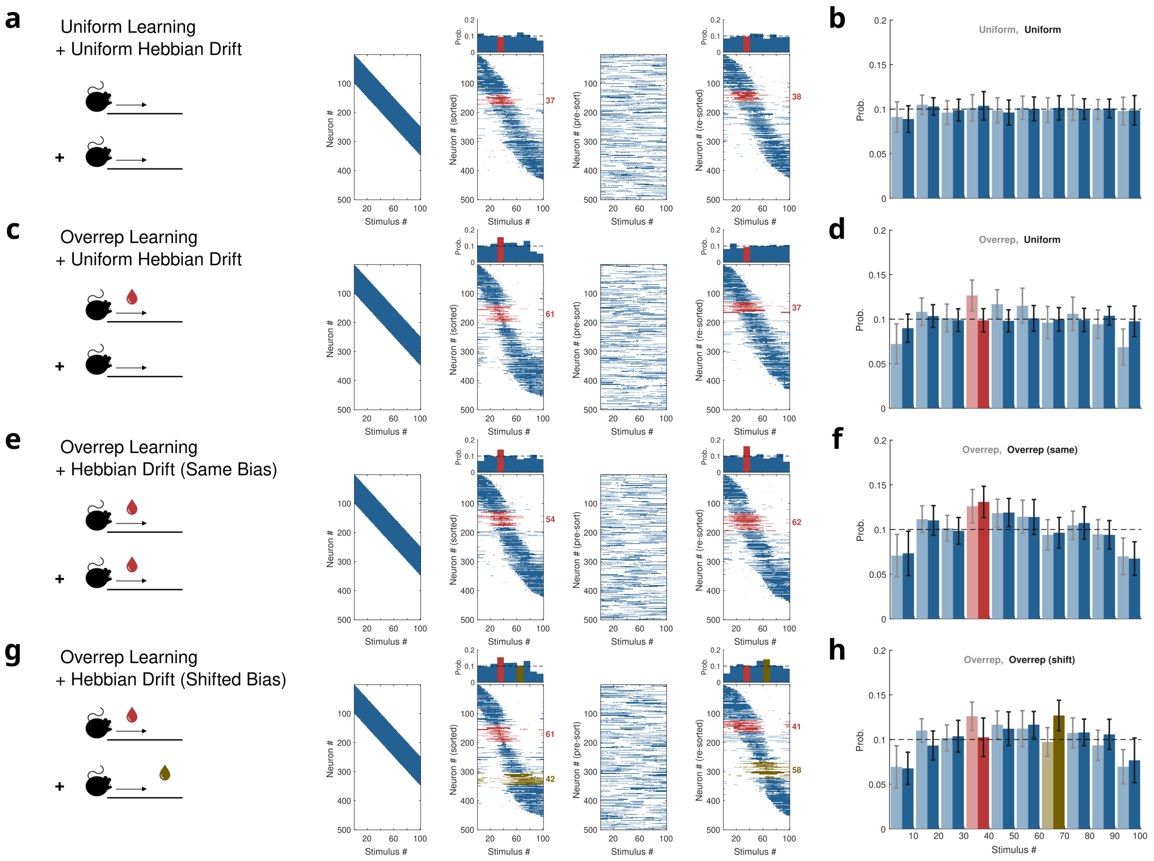

**Supplementary Fig. 6: Hebbian learning can preserve stimulus over-representation during drift. a)** Hebbian learning with random post-synaptic activity generates drifting network responses while preserving representational similarity between stimuli. **b)** Distribution of neurons with preferred stimulus in each 10-stimulus bin is flat after both stimulation periods. **c)** Over-representing stimuli during Hebbian updates (10-fold presentation frequency) creates over-representation in the activity space. This over-representation decays when switching to uniform stimulus presentation during subsequent Hebbian updates. **d)** Distribution of neurons with preferred stimulus in each 10-stimulus bin is elevated after initial over-representation but flat after uniform stimulation period. **e)** Drift preserves over-representation when the over-represented stimuli are maintained at higher presentation frequency throughout Hebbian updates. **f)** Distribution of neurons with preferred stimulus in each 10-stimulus bin is elevated both after initial over-representation and after second over-representation period with the same over-represeted stimulus. **g)** Changing which stimuli are over-represented during Hebbian updates correspondingly shifts the over-representation in activity space. **h)** Distribution of neurons with preferred stimulus in each 10-stimulus bin is elevated both after initial over-representation and after second over-representation period. The distribution follows the over-represented stimulus.

### Supplementary Methods

Here we provide additional analytical details supporting the main text. We performed all analytical calculations in terms of vector overlaps rather than angles; see Note 6 for the conversions used throughout.

We use the following notation: Input similarity is characterised by the **input overlap** $\gamma := \Pr(x_1^{(j)} = 1,\, x_2^{(j)} = 1)$, the joint activation probability of two input patterns at the same coordinate. Synaptic similarity between two networks $W_1$ and $W_2$ is characterised by the **synaptic overlap** $\beta := \Pr(W_1^{(ij)} = 1,\, W_2^{(ij)} = 1)$. Output angles are denoted $\alpha_{\mathrm{t}}$, $\alpha_{\mathrm{t+1}}$, and $\alpha_{\mathrm{drift}}$ as in the main text.

#### Note 1: Input-output mapping in a random binary feedforward network

Consider a feedforward network $\mathbf{y} = \sigma(W\mathbf{x})$, where $\mathbf{x} \in \{0,1\}^N$ is a sparse binary input with sparsity $f$, $W \in \{0,1\}^{M\times N}$ is a binary weight matrix with $K = gN$ non-zeros per row (network sparsity $g$), and $\sigma$ is a threshold nonlinearity enforcing output sparsity $f$. Inputs $\mathbf{x}_1, \mathbf{x}_2$ have sparsity $f$ and input overlap $\gamma$.

For a single output neuron $i$, the pre-activations $s_k = \sum_{j=1}^{N} W_{ij}\, x_k^{(j)}$ satisfy $\mathbb{E}[s_k] = Kf$ and $\mathrm{Var}(s_k) = Kf(1-f)$, with cross-covariance $\mathrm{Cov}(s_1, s_2) = K(\gamma - f^2)$. The pre-activation correlation is therefore

$$\rho = \frac{\gamma - f^2}{f(1-f)}, \tag{1}$$

which depends only on the input statistics $(f, \gamma)$ and is independent of both $K$ and $g$.

By the central limit theorem (large $K$), $(s_1, s_2)$ are approximately bivariate normal. Setting the threshold to the $(1-f)$-quantile of the Binomial$(K, f)$ distribution enforces output sparsity $f$, corresponding to the standard-normal cutoff $z_f = \Phi^{-1}(1-f)$. The output joint activation probability is

$$P_{11} = \Pr(Z_1 > z_f,\ Z_2 > z_f) = 1 - 2(1-f) + \Phi_2(z_f, z_f;\, \rho), \tag{2}$$

where $\Phi_2(\,\cdot\,; \rho)$ is the bivariate standard-normal CDF with correlation $\rho$. The output angle follows from $P_{11}$ via Note 6.

#### Note 2: Input–output similarity in general random networks

The result of Note 1 can be understood as a special case of a more general phenomenon. Consider a network layer $\mathbf{x} \mapsto \sigma\left(\frac{1}{\sqrt{n}} W\mathbf{x}\right)$, where $W \in \mathbb{R}^{n\times n}$ has i.i.d. entries and $\sigma$ is a pointwise nonlinearity. Let $\mathbf{x}_1, \mathbf{x}_2$ be mean-zero input vectors with i.i.d. components of variance $\sigma_x^2$ and pairwise covariance $\theta$. For large $n$, the pre-activations $\frac{1}{\sqrt{n}} w_{i\cdot}\mathbf{x}_k$ converge jointly to a bivariate Gaussian with covariance

$$\Sigma = \begin{pmatrix} \sigma_x^2 & \theta \\ \theta & \sigma_x^2 \end{pmatrix}. \tag{3}$$

The expected output similarity is

$$s_{\text{out}} = n\, \mathbb{E}_{z\sim\mathcal{N}(0,\Sigma)}\big[\sigma(z_1)\,\sigma(z_2)\big], \tag{4}$$

which is a monotonically increasing function of the input correlation $\theta/\sigma_x^2$. For the binary threshold nonlinearity ($\sigma = \mathbf{1}[\,\cdot > 0]$), this recovers Eq. (2) of Note 1.

**Singular-value analysis of representational distortion.** To characterise how a random network layer distorts the geometry of representations, consider the map $M = \frac{1}{\sqrt{m}}\sigma\Big(\frac{1}{\sqrt{n_{\text{in}}}}WZ\Big)$, where $W \in \mathbb{R}^{n_{\text{out}}\times n_{\text{in}}}$ and $Z \in \mathbb{R}^{n_{\text{in}}\times m}$ have i.i.d. standard Gaussian entries, with $m \gg n_{\text{in}}$ so that input directions are spread isotropically. Distortion is captured by the spread of the singular values of $M$: an isotropic singular spectrum implies that all input directions are stretched equally, while a spread spectrum implies direction-dependent amplification.

Using the linearisation $\sigma(x) \approx \zeta x + \sqrt{\rho^2 - \zeta^2}\,\varepsilon$ (where $\varepsilon$ is independent noise) together with operator-valued free-probability methods, the limiting singular spectrum is supported on $[\lambda_-, \lambda_+]$ with

$$\lambda_\pm = \sqrt{\big(\sqrt{\psi} \pm 1\big)^2 + \omega^2}, \qquad \omega^2 := \left(\frac{\rho}{\zeta}\right)^2 - 1 \;\geq 0, \tag{5}$$

where $\psi = n_{\text{out}}/n_{\text{in}}$ is the feature-expansion ratio, and

$$\rho := \sqrt{\mathbb{E}_z[\sigma^2(z)]}, \qquad \zeta := \mathbb{E}_z[\sigma'(z)]\,, \tag{6}$$

computed with respect to a standard normal $z$. The scalar $\omega^2$ measures the degree of nonlinearity of $\sigma$ ($\omega = 0$ for a purely linear layer).

A natural measure of distortion is the ratio of the largest to smallest singular value,

$$d = \sqrt{\frac{\big(\sqrt{\psi}+1\big)^2 + \omega^2}{\big(\sqrt{\psi}-1\big)^2 + \omega^2}}. \tag{7}$$

Three properties are worth noting. First, $d \to 1$ as $\psi \to \infty$: an expanding random layer introduces no representational distortion in the limit of large width, consistent with classical results on random projections. Second, $\omega^2 > 0$ (nonlinearity) keeps $d$ bounded even near $\psi = 1$, where a purely linear square random layer ($\omega = 0$) would diverge. Third, $d$ depends only on $\psi$ and $\omega$ and is independent of the specific weight realisation in the large-$n$ limit.

#### Note 3: Response vector drift

We consider a fixed input $\mathbf{x}$ presented to two networks $W_1, W_2$ with synaptic overlap $\beta$. For a single output neuron $i$, the pre-activations $s_k^{(i)} = \sum_j W_k^{(ij)} x^{(j)}$ satisfy $\mathbb{E}[s^{(i)}] = Ngf$ and $\mathrm{Var}(s^{(i)}) = Ngf(1-gf)$. Separating diagonal ($j = k$) and off-diagonal ($j \neq k$) contributions to the cross-moment gives

$$\mathrm{Cov}\Big(s_1^{(i)}, s_2^{(i)}\Big) = Nf(\beta - g^2 f), \tag{8}$$

so that the pre-activation correlation is

$$\rho = \frac{\beta - g^2 f}{g\,(1-gf)}. \tag{9}$$

Note that $N$ cancels. The output overlap $P_{11}$ and drift angle $\alpha_{\mathrm{drift}}$ follow via Eq. (2) and Note 6.

**Relation to rewiring fraction.** For a random network in which a fraction $d$ of connections is independently removed and redrawn, $\beta = (1-d)\,g$, giving

$$\rho = \frac{(1-d) - gf}{1-gf}, \tag{10}$$

which decreases monotonically from 1 at $d = 0$ to $-gf/(1-gf)$ at $d = 1$.

#### Note 4: Response angle drift

We now ask how the angle between responses to a fixed input pair $(\mathbf{x}_1, \mathbf{x}_2)$ changes as the network drifts from $W_1$ to $W_2$. The input overlap is $\gamma$ and the synaptic overlap is $\beta$.

**Variance of the similarity difference.** Because $W_1$ and $W_2$ share the same marginal statistics, the expected similarity difference $\mathbb{E}[s_1^{(i)} s_2^{(i)} - s_1'^{(i)} s_2'^{(i)}] = 0$. The effect of drift is therefore captured entirely by the variance. Expanding $\mathbb{E}\Big[(s_1^{(i)} s_2^{(i)} - s_1'^{(i)} s_2'^{(i)})^2\Big]$ and enumerating the distinct index configurations yields

$$\begin{aligned}\mathrm{Var}_{\mathrm{lin}}(\gamma,\beta) = {} & N\gamma\,(2g-2\beta)\\ & + N(N-1)\,\gamma f\cdot 2g\,(g-\beta)\\ & + N(N-1)\cdot 2f^2\,(g^2-\beta^2)\\ & + N(N-1)\cdot 2\gamma^2\,(g^2-\beta^2)\\ & + N(N-1)(N-2)\cdot 2f^3g^2\,(g-\beta)\\ & + N(N-1)(N-2)\cdot 2\gamma f^2g^2\,(g-\beta). \end{aligned} \tag{11}$$

**From pre-threshold variance to binarised outputs.** We map $\mathrm{Var}_{\mathrm{lin}}$ to an effective pre-activation correlation via

$$\rho_{\mathrm{lin}}(\gamma,\beta) = \frac{\mathrm{Var}_{\mathrm{lin}}(\gamma,\beta)}{\sigma_s^2+\mathrm{Var}_{\mathrm{lin}}(\gamma,\beta)}, \qquad \sigma_s^2 = Ngf(1-gf), \tag{12}$$

and compute $P_{11}(\gamma,\beta)$ via Eq. (2). Since the two networks are statistically independent, the variance of the population-mean cosine difference is

$$\mathrm{Var}_{\mathrm{bin}}(\gamma,\beta) = \frac{2}{N}\,P_{11}(\gamma,\beta)\big(1-P_{11}(\gamma,\beta)\big). \tag{13}$$

**From cosine fluctuations to angular deviation.** Linearising the angle–cosine mapping around the reference output angle $\alpha_0$, the expected absolute angular deviation is

$$\mathbb{E}[|\Delta\alpha|] \approx \frac{\sqrt{2/\pi}}{\sin(\alpha_0)}\sqrt{\mathrm{Var}_{\mathrm{bin}}(\gamma,\beta)}. \tag{14}$$

#### Note 5: Effects of output dimensionality on representational similarity

How does representational similarity depend on the dimensionality ratio between input and output spaces? The Johnson-Lindenstrauss lemma [**?**] establishes that random projections preserve pairwise distances in high-dimensional spaces; here we examine how smaller output dimensionality $N_{\mathrm{out}}$ affects this preservation when $N_{\mathrm{in}}$ and $N_{\mathrm{out}}$ are independent parameters.

For the **input-output mapping**, expectation values of the pre-activation statistics depend only on the fan-in $K = gN_{\mathrm{in}}$ and are independent of $N_{\mathrm{out}}$. The output angle relationship thus has the same expectation across all output sizes. However, random compression reduces the fidelity of response angles. Additionally, the binarization nonlinearity introduces quantization noise.

For **response vector drift**, the pre-activation correlation $\rho$ depends only on $g$, $f$, and synaptic overlap $\beta$, remaining independent of both $N_{\mathrm{in}}$ and $N_{\mathrm{out}}$. Expectation values are therefore unchanged. Again, the small output dimension leads to higher variance.

**Response angle drift**, however, depends on the variance, not the mean (which has expectation zero). The variance explicitly depends on $N_{\text{out}}$ (Eq. 13):

$$\mathrm{Var}_{\text{bin}}(\gamma,\beta) = \frac{2}{N_{\text{out}}} P_{11}(\gamma,\beta)[1 - P_{11}(\gamma,\beta)] \tag{15}$$

Thus response angle drift scales as $1/\sqrt{N_{\text{out}}}$, with leading-order behavior $\propto N_{\text{out}}^{-1/2}$.

While compression leaves the mean expected output and response vector drift unchanged, it increases the variance and thus directly affects response angle drift, our measure for representational similarity changes.

#### Note 6: Hebbian networks (derived in [1])

**Steady-state statistics.** Under the binary Hebbian rule with parameters $(p_+, p_-)$, the network converges to a steady-state connection probability

$$\bar{c} = \frac{f^2 p_+}{f^2 p_+ + 2f(1-f)p_-}, \tag{16}$$

with characteristic rate $\lambda = f^2 p_+ + 2f(1-f)p_-$. The Hebbian rule introduces correlations between synapses sharing the same postsynaptic neuron, characterised by the joint potentiation probability

$$\bar{c}_{11} = \frac{f^2 p_+[f(p_+ + 2(1-p_+)\bar{c}) + 2(1-f)(1-p_-)\bar{c}]}{1 - f[f(1-p_+) + (1-f)(1-p_-)]^2 - (1-f)[1 - fp_-]^2}. \tag{17}$$

After $T$ additional update steps at learning rate $\eta$, the synaptic overlap between the original and drifted network decays as

$$\beta = \bar{c}\, e^{-T\eta\lambda}. \tag{18}$$

**Modified pre-activation statistics.** The Hebbian rule modifies the pre-activation statistics through the inter-synapse correlation $\bar{c}_{11} - \bar{c}^2$. The pre-activation variance (common denominator for all settings) becomes

$$\mathrm{Var}(s) = fN\,\bar{c}(1-\bar{c}) + fN(fN-1)\,(\bar{c}_{11} - \bar{c}^2). \tag{19}$$

For the **input-output mapping** (two inputs, one network at steady state), the covariance numerator is

$$\mathrm{Cov}_{\text{in-out}} = \gamma N\,\bar{c}(1-\bar{c}) + (f^2N^2 - \gamma N)\,(\bar{c}_{11} - \bar{c}^2), \tag{20}$$

giving $\rho_{\text{Hebb}}^{\text{in-out}} = \mathrm{Cov}_{\text{in-out}}/\mathrm{Var}(s)$.

For **response vector drift** (one input, network at times $t_1$ and $t_2$), the off-diagonal inter-synapse correlation also decays with time, giving

$$\mathrm{Cov}_{\mathrm{vec}} = fN\big(\beta - \bar{c}^2\big) + (f^2N^2 - fN)\,(\bar{c}_{11} - \bar{c}^2)\,\frac{\beta}{\bar{c}}, \tag{21}$$

and $\rho^{\mathrm{vec}}_{\mathrm{Hebb}} = \mathrm{Cov}_{\mathrm{vec}}/\mathrm{Var}(s)$.

For **response angle drift**, the variance formula (11) applies with $g$ replaced by $\bar{c}$ and $\beta = \bar{c}\,e^{-T\eta\lambda}$ from Eq. (18).

In each case, the output overlap $P_{11}$ and all derived angles are computed via Eq. (2) and Note 6, exactly as for the random network.

#### Note 7: Conversion utilities

**Overlap to angle.** For binary patterns of sparsity $f$, exactly $\gamma N$ neurons are co-active out of $fN$ active neurons per pattern, so the cosine similarity equals $\gamma/f$. The angle and overlap are therefore related by

$$\alpha = \arccos\left(\frac{\gamma}{f}\right), \qquad \gamma = f\cos(\alpha). \tag{22}$$

**Overlap to Pearson correlation.** For binary outputs of sparsity $f$, the joint "on" probability $P_{11}$ and the Pearson correlation $\rho_{\mathrm{bin}}$ are related by the linear transform

$$\rho_{\mathrm{bin}} = \frac{P_{11} - f^2}{f(1-f)}, \qquad P_{11} = f(1-f)\,\rho_{\mathrm{bin}} + f^2. \tag{23}$$

**Synaptic overlap to rewiring fraction.** For the random network, a rewiring fraction $d$ gives $\beta = (1-d)\,g$ (see Note 3). For the Hebbian network, $T$ update steps at rate $\eta$ give $\beta = \bar{c}\,e^{-T\eta\lambda}$ (see Note 5).
